# Activation parameters, enthalpy-entropy compensation and the temperature-dependent activity of enzymes

**DOI:** 10.1101/2025.11.16.688712

**Authors:** Matthew J. McLeod, Robert E. Thorne

## Abstract

The increase in enzyme-catalyzed reaction rates with temperature is typically modeled using Arrhenius or Eyring relations. Interpretation of extracted parameters is subject to multiple caveats. Here we analyze the impact of expected small temperature variations of underlying activation/Eyring parameters on this modeling. Linear Arrhenius/Eyring behavior can still be observed when the underlying activation energy *E*_*a*_ or enthalpy Δ*H* and entropy Δ*S* vary with temperature. Modest variations — of the order of an H-bond energy over 60 °C — lead to large fractional deviations of *E*_*a*_, Δ*H* and Δ*S* values derived from linear fits from their underlying values and to deviations of Arrhenius prefactors *A* by orders of magnitude. In a family of related enzymes with similar activation free energies Δ*G*, small differences in temperature variation of Δ*H* and Δ*S* will lead to apparent enthalpy-entropy compensation and may scramble enzyme ordering based on Δ*H* or Δ*S*. For enzymes from cold and warm-adapted species having largely similar active sites, small temperature variations of Δ*H* and Δ*S* may explain large differences in apparent Δ*H* values. Similar considerations apply to interpretation of van ‘t Hoff plots of equilibrium measurements and related observations of enthalpy-entropy compensation. Complementary methods including simulations and multi-temperature static and time-resolved atomic-resolution structural studies should play a key role in interpreting temperature-dependent kinetic and equilibrium data from enzymatic systems.

**One-Sentence Summary:** Linear Arrhenius/Eyring behavior can occur even when enzyme structure and interactions vary with temperature, yielding fit parameters far from underlying values and generating apparent enthalpy-entropy compensation.

## 1. Introduction

Chemical reaction rates usually increase with temperature. For small-molecule catalysts, increased rates arise from increased collision velocities and frequencies. Observed reaction rates versus temperature are fit using Arrhenius (Arrhenius, 1889) or Eyring -Polanyi (Eyring, 1935c; Evans & Polanyi, 1935) (E-P) formalisms. The fit parameters — activation energy *E*_*a*_ and prefactor *A*, or enthalpy Δ*H* and entropy Δ*S* — are assumed to be independent of temperature within temperature ranges where the slope of Arrhenius or Eyring plots is constant.

Small-molecule catalysts are often structurally “locked down” — with tightly constrained conformational ensembles and largely temperature-independent interactions — and their Arrhenius plots may be linear over a wide temperature range. There, the assumption of temperature-independent activation parameters may be reasonable.

For enzyme-catalyzed reactions (Garcia-Viloca, Gao, Karplus, & Truhlar, 2004; Purich, 2010; Truhlar, 2015), analysis often begins with the kinetic rate coefficient *k*_*cat*_, which is measured under saturating substrate conditions. Turnover (*k*_*cat*_) then reports on steps after formation of the Michaelis complex and may reflect multiple steps. Interactions relevant to both enzyme structure and chemistry – hydrophobic, electrostatic, and hydrogen bonding – are temperature dependent and finely balanced, leading to small and large variations in structural ensembles and in function with temperature including changes in rate limiting steps (Machado, Gloster, & Da Silva, 2018).

Despite these and other significant mechanistic differences, when Arrhenius or Eyring-Polanyi (E-P) plots of enzymatic *k*_*cat*_ data are approximately linear over some (often small) temperature interval, the fit parameters Δ*H* and Δ*S* are again assumed to be temperature-independent, and the parameters so obtained may be used to compare distinct or related (e.g., mutant, thermally adapted) enzymes. For a family of related enzymes, a plot of the fit parameters Δ*S* versus Δ*H* often reveals a linear correlation with a correlation coefficient near 1 and may be interpreted as evidence for enthalpy-entropy compensation (EEC) (Chodera & Mobley, 2013; Cornish-Bowden, 2002; Exner, 1973; Fox, Zhao, Fink, Kang, & Whitesides, 2017; Liu & Guo, 2001). The interpretation of fit parameters derived from Arrhenius/E-P (and van ‘t Hoff) fits to enzymatic data is subject to multiple caveats, and the interpretation of apparent enthalpy-entropy compensation is controversial (Exner, 1973; Krug, Hunter, & Grieger, 1976b, 1976a; Cornish-Bowden, 2002; Fox et al., 2017; Barrie, 2012; Cornish-Bowden, 2017; Khrapunov, 2018; Griessen & Dam, 2021).

Theoretical analysis of enzymatic reactions using quantum mechanical/molecular mechanical (QM/MM) simulations, particularly using the empirical valence bond (EVB) method, have successfully predicted free energies of activation Δ*G*‡ and their temperature dependence for many enzymes (Åqvist & Brandsdal, 2025; Garcia-Viloca et al., 2004). As the corresponding entropy Δ*S*‡ is difficult to directly evaluate using these methods (Kazemi & Åqvist, 2015), Δ*H*‡ and Δ*S*‡ may be obtained from linear fits to plots of the calculated Δ*G*‡ vs *T*, again, with the assumption that both Δ*H*‡ and Δ*S*‡ are temperature independent (Åqvist & Brandsdal, 2025). However, even in a temperature interval where an enzymatic rate is determined solely by a single transition state ensemble, Δ*H*‡ and Δ*S*‡ (derived from partition functions of the activated complex and reactant states in transition state theory (Eyring, 1935a, 1935b; Hynes, Laage, Tuñón, & Moliner, 2016; Peters, 2017)) are expected to vary with temperature.

Recent multi-temperature near-atomic resolution structural studies of the metabolic enzyme phosphenolpyruvate carboxykinase (PEPCK) found that catalytically relevant structural changes occur in a temperature range where the Arrhenius/E-P plot shows an approximately linear variation, and where the Arrhenius/E-P fit parameters would normally be assumed to be independent of temperature (McLeod, Barwell, Holyoak, & Thorne, 2025). Motivated by this and other observations of how temperature affects structure (Keedy et al., 2018, 2015), we examined the impact of temperature-dependent parameters on Arrhenius/E-P plots and on fit parameters obtained from those plots.

## 2. Results

### 2.1 Arrhenius fits and anomalously large prefactors

In the Arrhenius model for chemical reaction rates (Arrhenius, 1889),

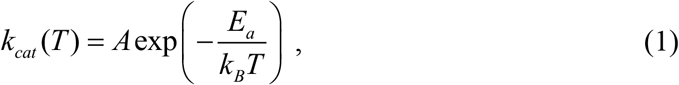

Where *E*_*a*_ is the activation energy and *A* is a prefactor. Fits of plots of ln (*k*_*cat*_) vs 1/*T* to

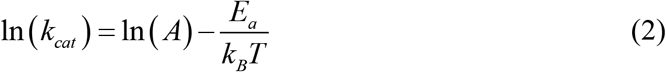

are used to extract empirical parameters *E*_*a*_ and *A* from the slope and intercept of the fit, respectively. This implicitly assumes that both *E*_*a*_ and *A* are independent of temperature over the range of the fit.

In the following, we will distinguish between the “apparent” parameters 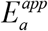 and *A*^*app*^ derived from a linear fit to data on Arrhenius axes and the “underlying” parameters *E*_*a*_ (*T*) and *A*(*T*) that, e.g., might be derived from an accurate first-principles calculation or from measurements that directly probed each parameter.

Suppose that *E*_*a*_ (*T*) varies linearly with temperature about some temperature *T*_0_ within the measured /biological temperature range,

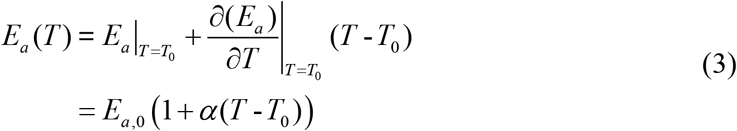

Where

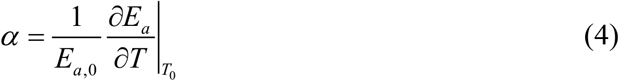

is the fractional rate of change of *E*_*a*_ with temperature near *T*_0_. Formally, Eq. 3 corresponds to the first two terms in a Taylor series expansion of *E*_*a*_ (*T*) about *T*_0_ and so provides a general description. Then

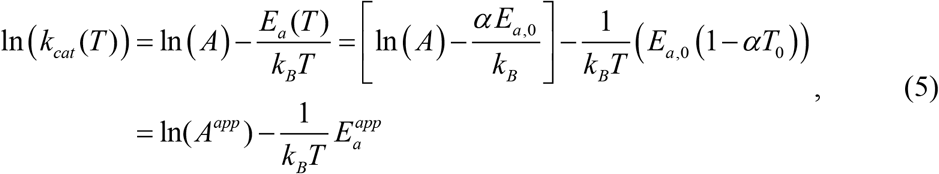

so that *k*_*cat*_ (*T*) can be written as

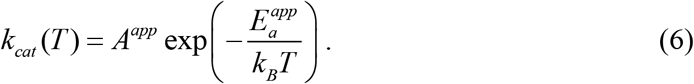

The resulting Arrhenius plot will still be strictly linear. The slope and 1/*T* = 0 K^-1^ intercept will be set by

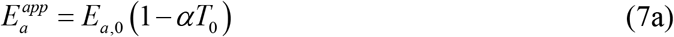

and

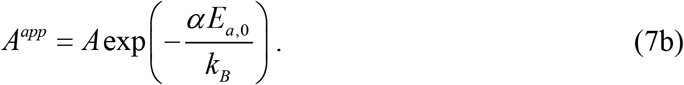

Taking the log of Eq. 7b and using Eq. 7a gives

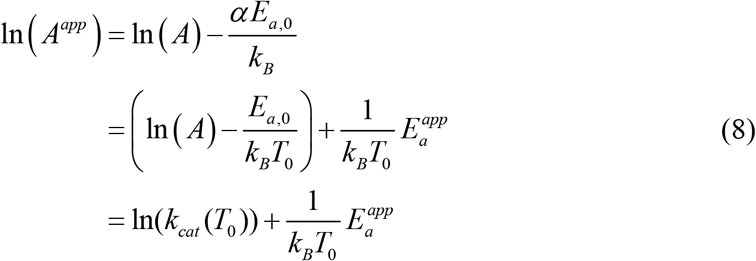

Equations 7-8 indicate four consequences of a linear temperature variation of *E*_*a*_. First, the *E* apparent activation energy 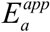 derived from the Arrhenius fit is changed relative to *E*_*a*,0_ = *E*_*a*_ (*T* = *T*_0_) by the fractional rate of change of *E*_*a*_ (*T*) with temperature, *α*, times the absolute temperature *T*_0_.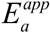 is equal to the value *E*_*a*_ would have at *T* = 0 K if its linear temperature *E* variation near *T* = *T*_0_ was maintained to absolute zero. If the activation energy *E*_*a*_ (*T*) decreases *E* with increasing temperature (i.e., *α* < 0), the apparent activation energy 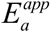 will be larger than *E*_*a*_ (*T*_0_). **Table S1** gives the apparent activation energy 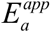 and prefactor *A*^*app*^ assuming a linear variation of *E*_*a*_ about *T*_0_=303 K for several values of the temperature coefficient *α*. **Figure S1** shows *E*_*a*_ vs *T* and the resulting Arrhenius plot in each case.

Typical activation energies for enzymatic reactions range from ∼2 to ∼35 kcal/mol or ∼1,000-17,500 °Cin temperature units. Over a biological temperature range, e.g., from 0°C to 60°C, it is not implausible that *E*_*a*_ (*T*) could decrease by 10%, corresponding to a temperature coefficient *α=*−0.167%/°C, due to small shifts in, e.g., active site/substrate conformation, interaction strengths, or solvent structure. A negative temperature coefficient of this magnitude will increase the apparent activation energy — deduced from the slope of the Arrhenius plot — by a multiplicative factor of 1.51 relative to *E*_*a*,0_. For an activation energy of 10 kcal/mol, a 10% change in activation energy over 60 °C is only 1 kcal/mol, comparable to the energy required to lengthen a single hydrogen bond by 0.1 Å (Herschlag & Pinney, 2018); the apparent activation energy is increased by 5.1 times this amount.

A second consequence of temperature variation of *E*_*a*_ is that *A*^*app*^ /*A*, the ratio of apparent and actual prefactors, is exp(−*α E*_*a*,0_ /*k*_*B*_). For *α=*−0.167%/°C and activation energies of 10 kcal/mol and 35 kcal/mol, *A*^*app*^ /*A* is 4.5 × 10^3^ and 6.0 × 10^12^, respectively. Large prefactors are often obtained from Arrhenius fits to data for enzymatic activity, protein denaturation (Qin, Balasubramanian, Wolkers, Pearce, & Bischof, 2014), and other biophysical data (Liu & Guo, 2001), far larger than may be reasonably accounted for by “loose” (i.e., high entropy) transition states (Robinson, P. J. & Holbrook, K. A., 1972). Small fractional decreases of *E*_*a*_ with temperature provide one possible explanation for large prefactors.

A third consequence of temperature variation of *E*_*a*_ is that neither the underlying *E*_*a*_ (*T*) nor the underlying Arrhenius prefactor *A* can be determined from the Arrhenius fit. The linear fit and its derived parameters *A*^*app*^ and 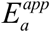 provide a constraint on the values and temperature dependences of the underlying parameters, but they do not determine them.

A fourth consequence of a temperature-dependent *E*_*a*_ is a mechanism for generating correlation between the fit parameters *A*^*app*^ and 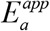. Consider a series of enzymes whose linear Arrhenius fits give similar values of *k*_*cat*_ (*T*_*c*_) at some temperature *T*_*c*_, and whose activation energies *E*_*a*_ (*T*) vary linearly with temperature with different coefficients *α*. We can take *T*_0_ = *T*_*c*_, so that Eq. 8 then has the form

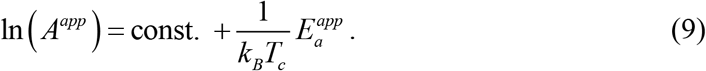

Consequently, for such a series of enzymes there will be a linear correlation between the log of the apparent pre-exponential factor *A*^*app*^ and the apparent activation energy 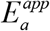. Such a correlation has been observed for a variety of systems and often involves unphysically tight correlations.

Note that we have made the simplifying assumption that the prefactor *A* is temperature independent. Relaxing this assumption for *A* does not change the major conclusions, as implied by the following analysis.

### 2.2 Eyring fits and enthalpy-entropy compensation

In the transition state model, initially presented by Eyring and Polanyi (Eyring, 1935b; Evans & Polanyi, 1935),

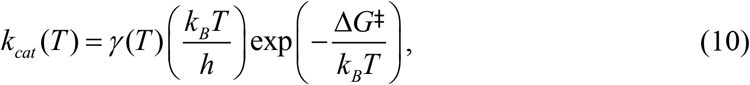

Where Δ*G*‡ = Δ*H* ‡ − *T* Δ*S*‡ is the free-energy difference between the ground state (GS) and transition state (TS) (the activation free energy) and *γ* (*T*) is the transmission coefficient. For simplicity, we ignore contributions from dividing surface recrossing, tunneling and non-equilibrium effects (Garcia-Viloca et al., 2004) and assume *γ* (*T*) =1, as is common when fitting experimental data. Parameters of an Arrhenius fit (Eq. 1) are then given by

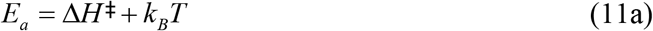

and

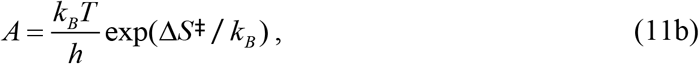

where the second term in Eq. 11(a) can usually be neglected.

The Eyring-Polanyi model applies in its derived form only when *k*_*cat*_ (*T*) is solely determined by a single transition state (or dividing surface). In enzymes, *k*_*cat*_ (*T*) is determined by all steps forward of enzyme-substrate (ES) complex formation. This may include multiple chemical steps, including more than one step that contributes substantially to the observed rate. This also includes non-chemical steps such conformational changes required for successive chemical steps and for product release, known to be rate limiting in many enzymes. Consequently, *k*_*cat*_ (*T*) may or may not be dominated by a single transition state in the experimental temperature range, and the relative contributions of different steps to the overall rate may vary over that range. The application of Eq. 10 with *γ* (*T*) =1 as a fit to enzyme data to extract enthalpy and entropy changes is thus in general phenomenological.

To describe experimental data, Eq. 10 can be rewritten (with *γ* (*T*) =1) as

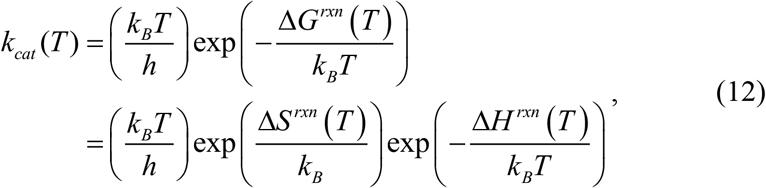

where the “rxn” labeling now indicates that the quantities parameterizing the reaction may not be solely determined by a transition state /dividing surface. If Δ*H*^*rxn*^ and Δ*S*^*rxn*^ are both independent of temperature, the slope and 1/*T* = 0 K intercept of a linear fit to a plot of log (*k*_*cat*_ (*T*) *T*) vs 1/*T* then give Δ*H*^*rxn*^ and Δ*S*^*rxn*^, respectively.

We again distinguish between the underlying values Δ*H* ^*rxn*^ (*T*) and Δ*S*^*rxn*^ (*T*), as might be obtained from an accurate first principles calculation, and the apparent temperature-independent values Δ*H*^*rxn,app*^ and Δ*S*^*rxn,app*^ obtained from the slope and intercept of a linear fit to an Eyring plot.

Generalizing our Arrhenius analysis of Section 2.1, assume the underlying Δ*G*^*rxn*^ (*T*) varies linearly about some temperature *T*_0_ in the biological temperature range so that

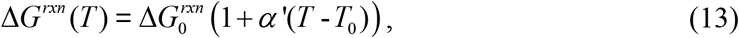

where 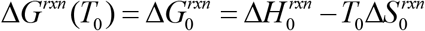 and

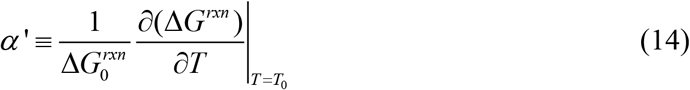

is the fractional rate of change of Δ*G*^*rxn*^ (*T*) with temperature (e.g., in %/°C). Eq. 12 can be written as

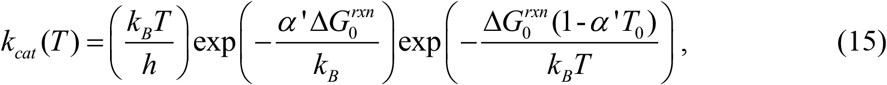

so that ln(*k*_*cat*_ /*T*) has the form

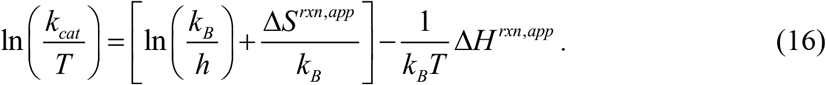

The slope and 1/*T* = 0 K intercept of an Eyring plot give

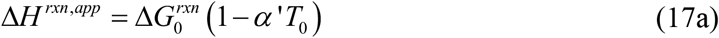

and

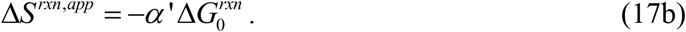

When 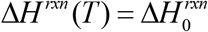 and 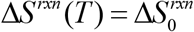 are independent of temperature, Eqs. 17a and 17b reduce to 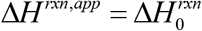 and 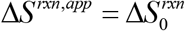, as required, and 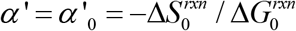 Because entropy is multiplied by −*T* in the Gibbs free energy, for temperature-independent Δ*H*^*rxn*^ and Δ*S* ^*rxn*^ ≠ 0, Δ*G*^*rxn*^ will still vary linearly with *T* and *α* ‘ ≠ 0.

These equations have multiple consequences. First, a linear Eyring plot requires only that Δ*G*^*rxn*^ (*T*) varies linearly with *T*. It does not imply that Δ*H*^*rxn*^ and Δ*S*^*rxn*^ are independent of temperature.

Second, knowledge of Δ*G*^*rxn*^ (*T*) (set by 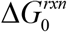 and *α* ‘) constrains but does not determine Δ*H* ^*rxn*^ (*T*) and Δ*S*^*rxn*^ (*T*). This is illustrated in **Fig. 1** and also in **Fig. S2**, which shows examples of Δ*H* (*T*), Δ*S* (*T*) pairs that are consistent with the given linearly-varying Δ*G*(*T*). Only if Δ*H* ^*rxn*^ (*T*) and Δ*S*^*rxn*^ (*T*) are assumed to be independent of temperature is there a unique solution given by 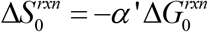 and 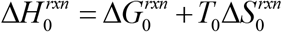.

**Figure 1.**
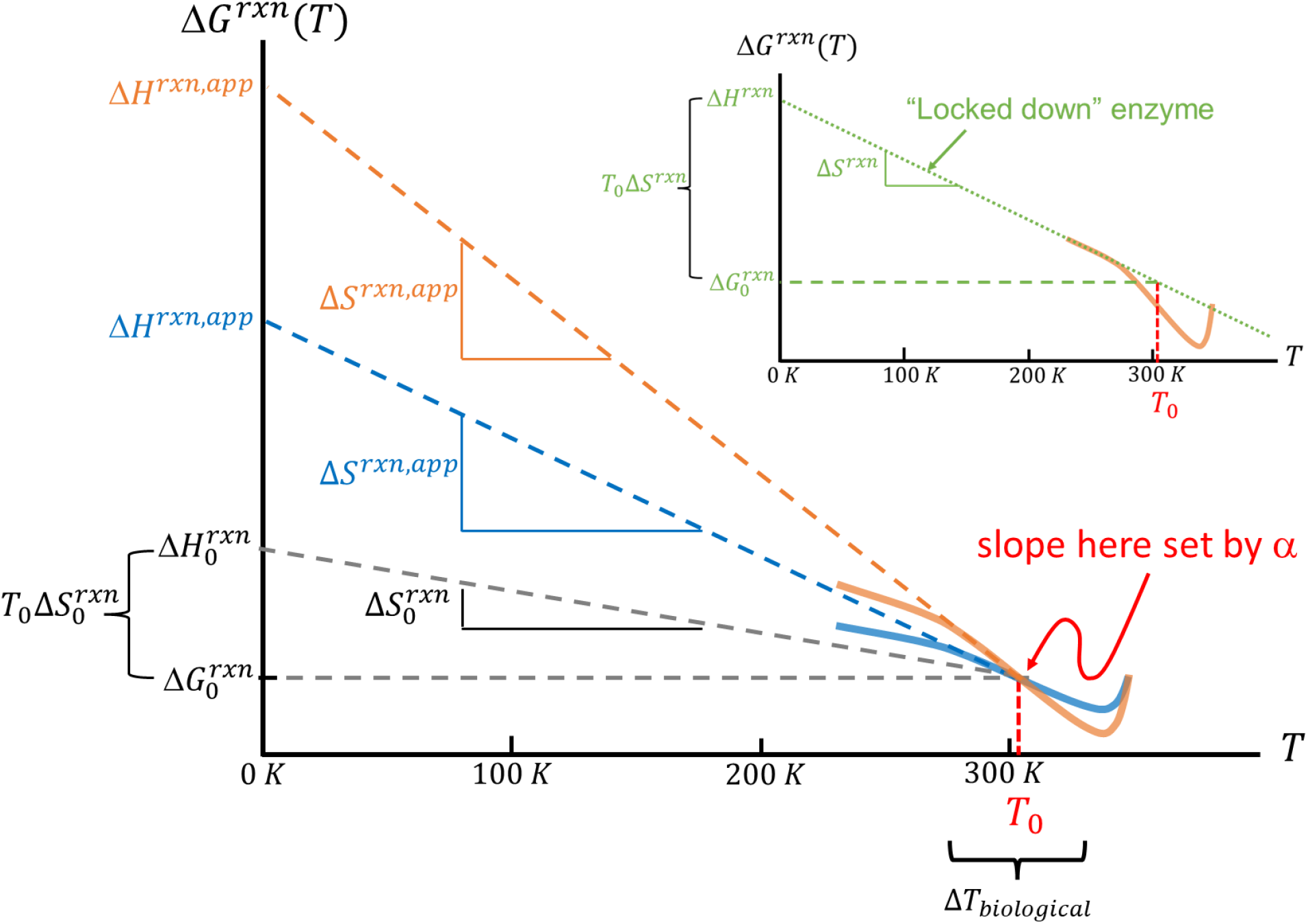
An Eyring plot of *k*_*cat*_ /*T* versus 1000/*T* gives Δ*G*^*rxn*^ (*T*). **Inset:** For a “locked down” enzyme, the slope of Δ*G*^*rxn*^ (*T*) would be roughly constant and equal to its low-temperature value. Conformational changes favorable (unfavorable) to catalysis in the real enzyme complex may increase (decrease) the reaction rate faster and thus decrease Δ*G*^*rxn*^ (*T*) faster (slower) in the biological temperature range than would occur without those changes. **Main panel:** If Δ*H*^*rxn*^ and/or Δ*S*^*rxn*^ varies with temperature (as assumed for the orange and blue curves), the Eyring fit parameters Δ*H*^*rxn,app*^ and Δ*S*^*rxn,app*^ (equal to the low-temperature intercept and slope of a linear fit to Δ*G*^*rxn*^ (*T*)) may show large deviations from the underlying values, even though the variations of Δ*H*^*rxn*^ and Δ*S*^*rxn*^ over the biological temperature range may be small.

Third, variations in the slope *α* ‘ of Δ*G*^*rxn*^ (*T*) near *T* = *T*_0_ will cause proportional variations in apparent enthalpy and entropy. The mathematics here is illustrated in **Fig. 2**. 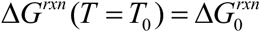 is determined by the enthalpy 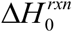 and entropy 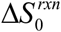 at that temperature. The slope of Δ*G*^*rxn*^ (*T*) near *T* = *T*_0_ (parameterized by 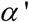) is determined by 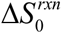 and by how Δ*H*^*rxn*^ and Δ*S* ^*rxn*^ are *changing*.

**Figure 2.**
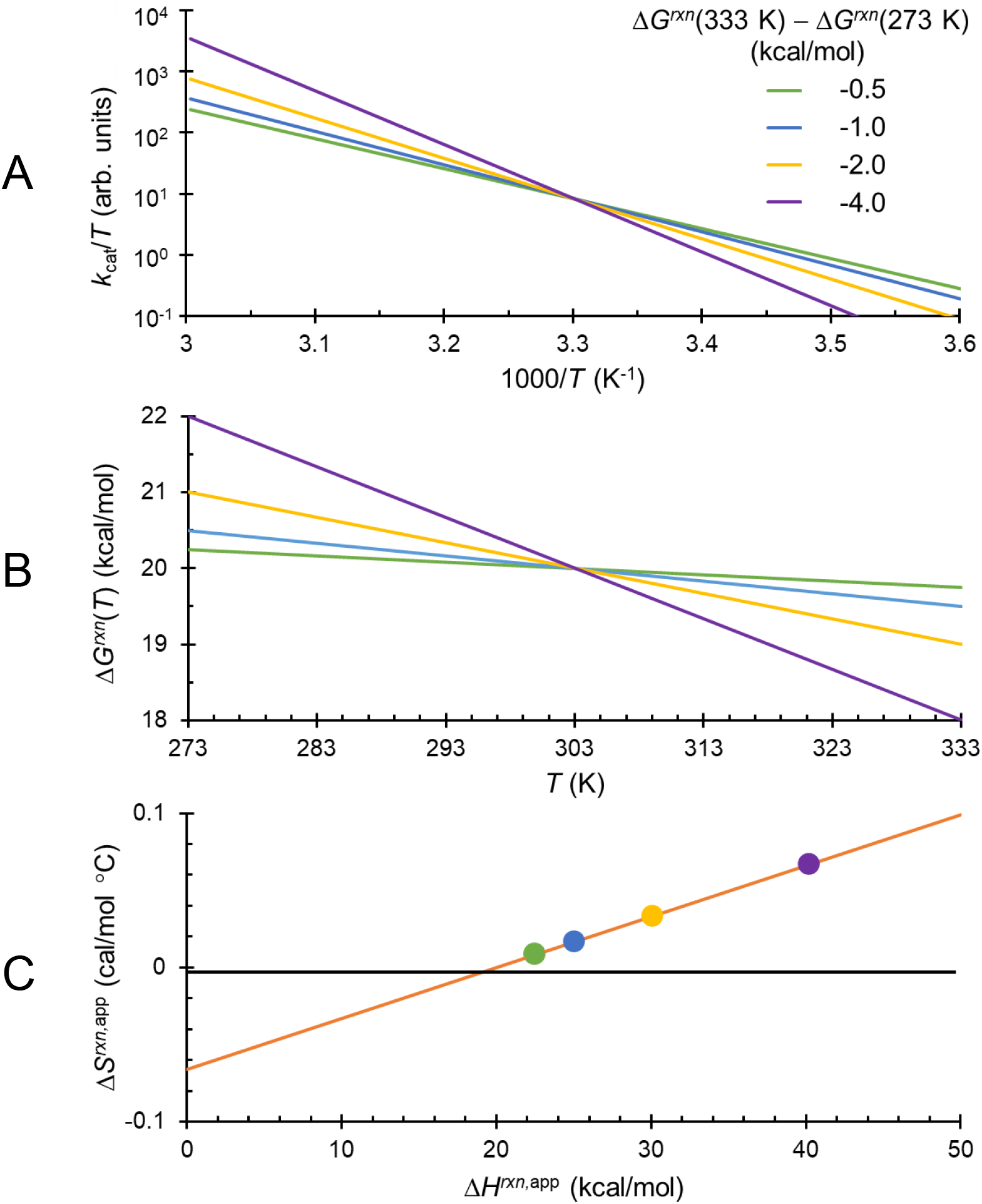
Underlying values of Δ*H*^*rxn*^ and Δ*S*^*rxn*^ and their temperature dependence cannot be determined from a linear fit to an Eyring plot. The fit only provides a constraint that must be satisfied at each temperature. **A** Linear Eyring plots with different slopes and the same *k*_*cat*_ /*T* values at *T* = *T*_0_ = 303 K. **B** The curves in **A** are calculated assuming Δ*G*^*rxn*^ (*T*_0_ = 303K) = 20 kcal/mol and temperature coefficients *α* of -0.04, -0.08, -0.17 and -0.33 kcal/mol °C. **C** Solid circles: Apparent enthalpy *H*^*rxn,app*^ and entropy *S* ^*rxn,app*^ deduced from the slope and intercept of each curve in the Eyring plot **A**, based on the assumption that Δ*H*^*rxn,app*^ andΔ*S*^*rxn,app*^ are independent of temperature. Δ*H*^*rxn,app*^ and Δ*S*^*rxn,app*^ values for all possible slopes of the Eyring plots with the same *k*_*cat*_ /*T* at *T* = *T*_0_ will fall on the orange line. The underlying values of Δ*H*^*rxn*^ and Δ*S*^*rxn*^ at *T* = *T*_0_ for each slope will also fall on the orange line but will otherwise be undetermined.

Fourth, the (temperature independent) apparent enthalpy and apparent entropy as deduced from a linear Eyring fit are related by

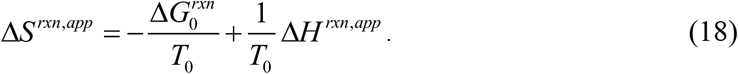

Consider a series of enzymes whose linear Eyring fits give similar values of *k*_*cat*_ (*T*_*c*_) and thus Δ*G*^*rxn*^ (*T*_*c*_) at some temperature *T*_*c*_, and whose free energies Δ*G*^*rxn*^ (*T*) vary linearly with temperature with different coefficients *α*’. We can take *T*_0_ = *T*_*c*_, so that Eq. 18 then has the form

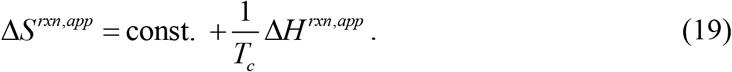

Consequently, the fit parameters Δ*S*^*rxn,app*^ and Δ*H*^*rxn,app*^ will be linearly correlated: there will be enthalpy-entropy compensation (**Fig. 3)** with a “compensation temperature” *T*_*C*_. More generally, the Eyring fits for the different enzymes will not cross at a single point (i.e., 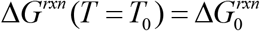 will not be the same for all enzymes.) In that case, *T*_*c*_ of a linear enthalpy-entropy fit will equal the temperature at which the variance of the Δ*G*^*rxn*^ (*T*) values for the different enzymes is minimized (Griessen & Dam, 2021), with the correlation coefficient of the linear fit – measuring the extent of scatter off the enthalpy-entropy fit – increasing with decreasing variance.

**Figure 3.**
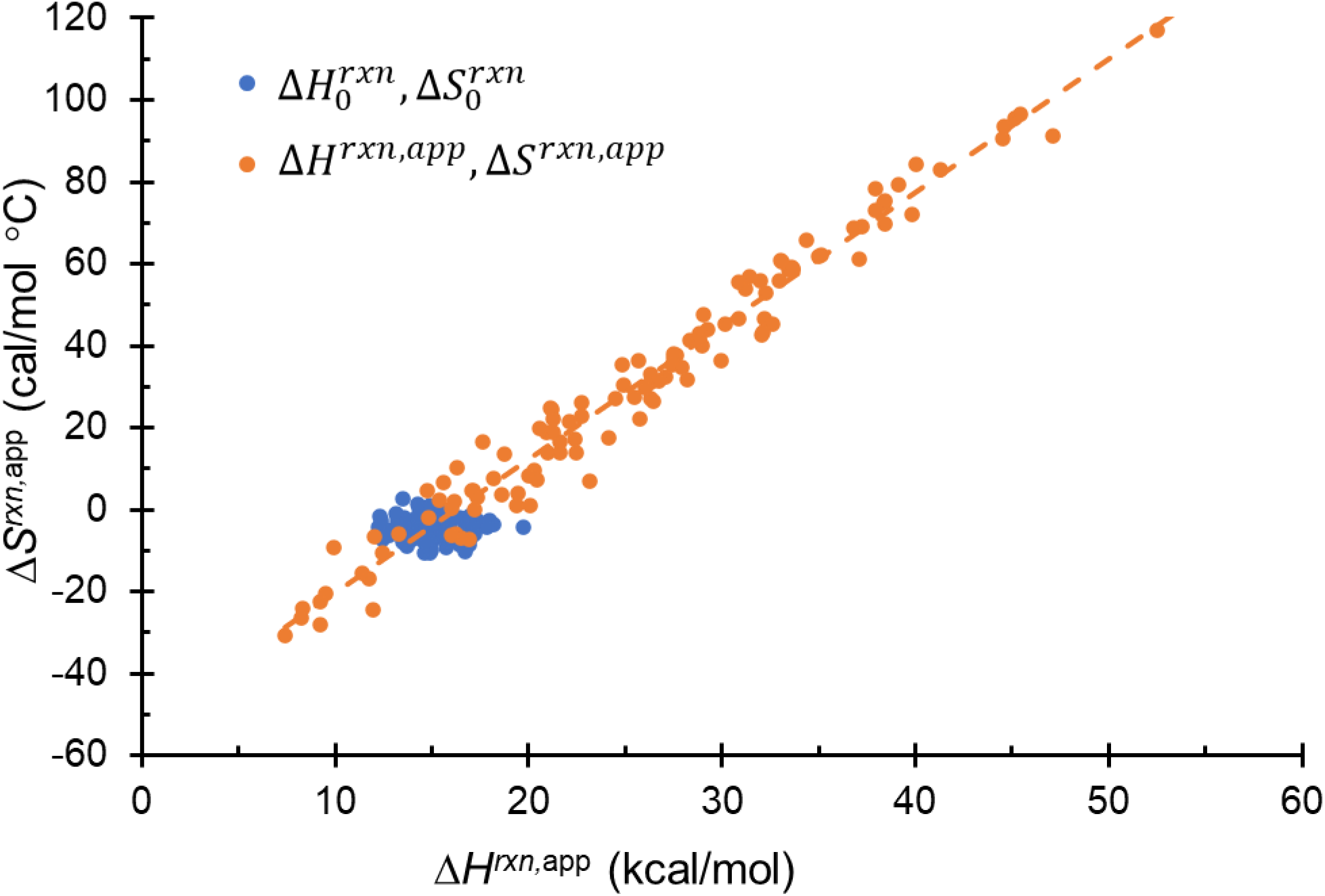
Eyring fit parameters may exhibit entropy-entropy compensation when the underlying enthalpy and entropy are not independent of temperature. **Blue symbols:** Randomly selected 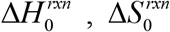 pairs, assuming that 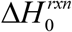 has a Gaussian distribution about 15 kcal/mol with a standard deviation of 1.5 kcal/mol, 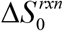 has a Gaussian distribution about -5 cal/mol °C with a standard deviation of 2.5 cal/mol °C, and the temperature coefficient *α*’ of Δ*G*^*rxn*^ (*T*) has a Gaussian distribution about -0.2 %/°C with a standard deviation of 0.2 %/°C. Δ*G*^*rxn*^ (*T*) and *k*_*cat*_ (*T*) are then calculated using Eqs. 12 and 13 with *T*_0_ = 303 K, for each randomly selected 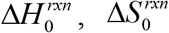, and *α*’. **Orange symbols:** Corresponding apparent enthalpy and entropy, Δ*H*^*rxn,app*^ and Δ*S*^*rxn,app*^, that would be extracted from a standard linear Eyring fit. These values show a strong linear dispersion /correlation. The free energy Δ*G*^*rxn*^ (*T* = *T*_0_) has a mean of ∼16.5 kcal/mol and a standard deviation of 1.7 kcal/mol. Small fractional variations in the temperature coefficient *α* lead to large fractional variations in apparent enthalpy and entropy and to apparent enthalpy-entropy compensation, even when the underlying enthalpy and entropy show no correlation/compensation.

### 2.3 Application to interpretation of kinetics measurements

As a first example, Fig. 3 of (McLeod et al., 2025) shows an Eyring plot of data for the dephosphorylation and carboxylation of phosphenolpyruvate (PEP) by the metabolic enzyme rat cytosolic PEPCK. An Eyring fit to the linear region between 8°C (281 K) and 25°C (298 K) gives Δ*H* ^*rxn,app*^ = 26.3 ± 0.85 kcal/mol and Δ*S*^*rxn,app*^ = 0.037 ± 0.003 kcal/mol•°C (*R*^2^=0.997), so that 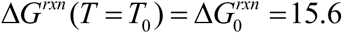 kcal/mol and *α* ‘ =0.24%/°C (∼0.04 kcal/mol•°C). The connection between the apparent values and the actual enthalpy Δ*H* ^*rxn*^ (*T*) and entropy Δ*S*^*rxn*^ (*T*) is unknown: the actual values need only satisfy 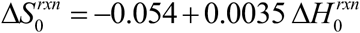 at *T* = *T*_0_ and sum to give Δ*G*^*rxn*^ (*T*) at other temperatures to be consistent with the Eyring fit.

As a second example, kinetic measurements of hydride transfer in WT and mutant thermophilic alcohol dehydrogenases (ht-ADH) indicated a break in Arrhenius slope at ∽30 °C (**Figures 4A, S3**) (Nagel, Dong, Bahnson, & Klinman, 2011). Above this temperature, all variants showed similar slopes and intercepts, and fits yielded physically reasonable Arrhenius prefactors *A*. Below this temperature, slopes varied substantially between variants and fits yielded prefactors as large as 10^25^ s^-1^. Large Arrhenius prefactors are not *a priori* unreasonable: from Eq. 11b they are expected when the process being fit involves a large positive change in entropy, e.g., in the case of a “loose” transition state (Baklanov & Kiselev, 2023; Robinson, P. J. & Holbrook, K. A., 1972). However, as shown in **Fig. 4D**, the large jumps in *A* are accompanied by large jumps in *E*_a_ such that there is near perfect compensation of the changes in ln(*A*) and *E*_a_.

**Figure 4.**
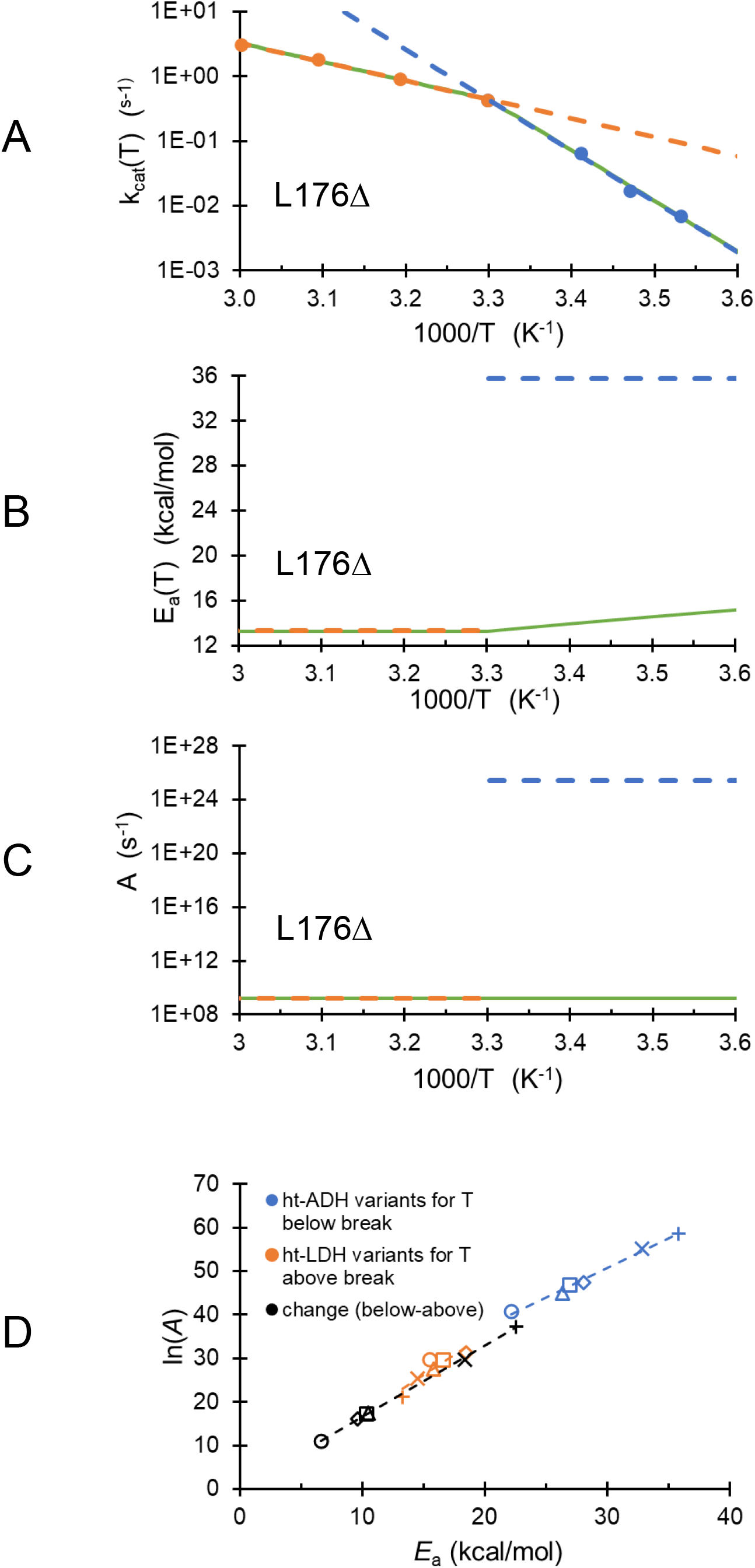
**A** Arrhenius plot of *k*_cat_(*T*) data from (Nagel et al., 2011) for a mutant form of a thermophilic alcohol dehdrogenase (ht-ADH). The orange and blue dashed lines are standard Arrhenius fits to the data above and below the break temperature, respectively. The corresponding apparent *E*_a_ and *A* values, determined by the slope and intercept of each fit, are shown by the orange and blue dashed lines in **B** and **C.** The dashed green line in **A** is a fit to the data assuming that *E*_a_ is constant above the break and varies linearly with temperature below the break (green line in **B**), and that *A* has the same value above and below the break (green line in **C**). **D** Plot of apparent ln(*A*) vs apparent *E*_a_ above (orange) and below (blue) the break obtained from Arrhenius fits to six ht-ADH variants. Above the break, ln(*A*) and *E*_a_ show some correlation, with a linear fit giving *R*^2^=0.82. Below the break, the correlation is stronger with *R*^2^=0.992. The black symbols indicate the net change in ln(*A*) and *E*_a_ that occurs at the break. These changes show a linear correlation with *R*^2^=0.9999.

**Table S2** gives the parameters Δ*H*^*rxn,app*^ and *T* Δ*S*^*rxn,app*^ obtained from linear Eyring fits, from Tables 2 and 3 in Ref. 31. **Figure S4** plots Δ*H*^*rxn,app*^ vs *T* Δ*S* ^*rxn,app*^ above and below the break for each variant, as well as the change in these values at the break. The changes in Δ*H*^*rxn,app*^ and *T* Δ*S* ^*rxn,app*^ at the break are nearly perfectly compensated. These results were accounted for by a model (**SI Section S1**) in which the enzyme populates only competent conformers above the break, and interconverts between competent and incompetent conformers below the break. Fits using this model required a free energy difference between the competent and incompetent conformers below the break of only 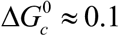 kcal/mol for all variants (**Table S3**).

One alternative starting point based on the Arrhenius model is to make no assumptions about mechanism other than to allow *E*_*a*_ (*T*) to vary with temperature. Since the prefactors are “reasonable” above the break at 30°C and “unreasonable” below, assume that *E*_*a*_ (*T*) is temperature-independent above the break and varies linearly with temperature below. **Table S4** gives the required temperature coefficient *α* and the corresponding change in *E*_*a*_ (*T*) between 0 and 30 °C required to replicate the measured Arrhenius slope and intercept for each enzyme variant, assuming *A* is fixed at its high temperature value. **Figure 4A** shows the resulting fit for one variant and **Figures 4B** and **4C** show *E*_*a*_ (*T*) and *A*(*T*); **Figure S4** shows corresponding plots for the other variants. In all cases, the required fractional variations of *E*_*a*_ (*T*) below the break are small and much smaller than the jumps in apparent *E*_*a*_ at the break.

Similarly, using the Eyring-Polanyi model, Δ*G*^*rxn*^ (*T*) can be assumed to vary linearly with temperature below and above the break at 30°C, with different slopes. Assuming that Δ*H* ^*rxn*^ (*T*) and Δ*S*^*rxn*^ (*T*) are constant above the break (so that an E-P fit directly gives their values and 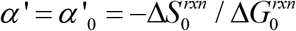 there), **Table S5** gives the required temperature coefficient *α* ‘ below the break to reproduce the fit E-P fit values Δ*H*^*rxn,app*^ and Δ*S*^*rxn,app*^ for each enzyme variant. The actual enthalpy and entropy values below the break cannot be determined and are constrained to lie upon a line given by Gibbs relation.

These alternative interpretations make few assumptions, yield reasonable parameters, and do not require precise cancellation of the enthalpic and entropic changes in the free energy above and below the break temperature for all mutants. The mechanism proposed in (Nagel et al., 2011) may be correct but other mechanisms may give the same behavior.

### 2.4 Application to thermal adaptation of enzymes

Eyring fits (assuming temperature-independent parameters) to kinetics measurements on enzymes from species adapted to different thermal environments indicate that psychrophilic enzymes generally have much lower Δ*H*^*rxn,app*^ and more negative Δ*S*^*rxn,app*^ values than their thermophilic and mesophilic counterparts, even though their activation free energies Δ*G*^*rxn*^ (*T*) near room temperature are usually similar (so there is enthalpy-entropy compensation) (Åqvist & Brandsdal, 2025; Feller & Gerday, 2003; Siddiqui & Cavicchioli, 2006). This difference has been suggested to arise from increased active site flexibility in psychrophilic variants (Fields & Somero, 1998). However, many such variants show largely identical active site structure and flexibility to those of mesophilic variants (Pedersen, Willassen, & Leiros, 2009; Smalås, Heimstad, Hordvik, Willassen, & Male, 1994; Tsigos et al., 2001). The differences in enthalpy and entropy have instead been attributed to the presence of flexible surface loops in psychrophilic enzymes (Åqvist & Brandsdal, 2025).

Molecular dynamics /empirical valence bond (MD/EVB) methods have quantitatively predicted the temperature-dependent free-energy of activation Δ*G*‡(*T*) for related thermally-adapted enzymes (Åqvist & Brandsdal, 2025). As the entropy Δ*S*‡(*T*) is difficult to evaluate using MD/EVB methods, the calculated Δ*G*‡(*T*) is fit assuming Δ*H*‡ and Δ*S*‡ are temperature-independent (as is also assumed in fitting the experimental data) (Åqvist & Brandsdal, 2025; Åqvist, Isaksen, & Brandsdal, 2017). MD/EVB calculations show that restraining surface loops located >18 Å away from the active site in the psychrophilic enzyme leads to a doubling of the enthalpy change, matching that of a mesophilic variant, with little effect on Δ*G*‡(*T*) (Isaksen, Åqvist, & Brandsdal, 2014, 2016).

**Figure 5A** shows activation free energies Δ*G*‡(*T*) calculated using MD/EVB versus inverse temperature for warm-adapted (bovine) and cold-adapted (arctic salmon) trypsins taken from Fig. 1 of (Isaksen et al., 2014), which closely agree with experimental Δ*G*^*rxn*^ (*T*) from Eyring fits. The orange and blue lines are fits to the Gibbs relation assuming the enthalpy and entropy changes are temperature independent, and give Δ*H* ‡ = 20.4 kcal/mol, Δ*S*‡ = +3.5 cal/mol °C and Δ*H* ‡ = 9.9 kcal/mol, Δ*S*‡ = −27.5 cal/mol °C, respectively.

**Figure 5.**
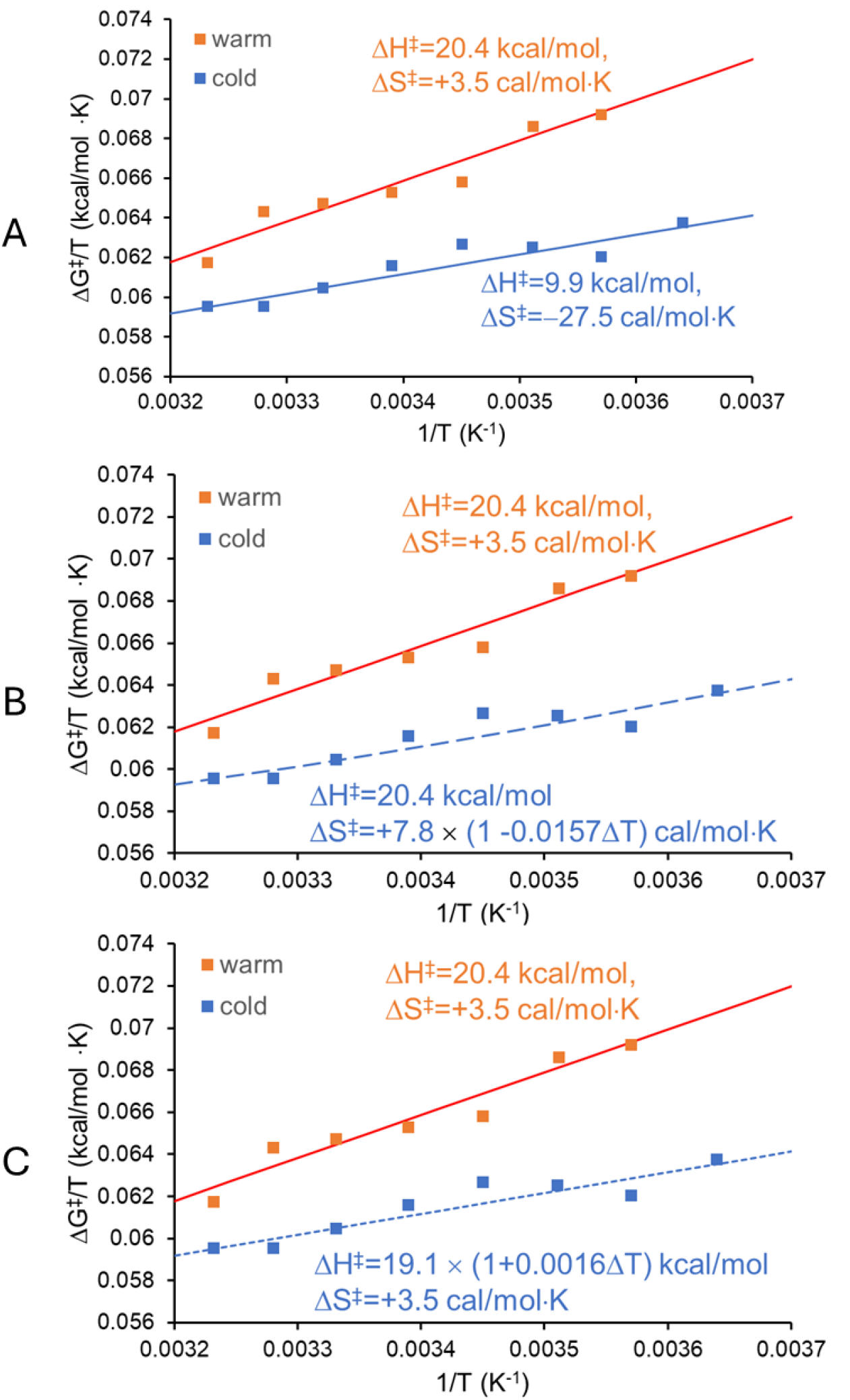
Symbols: Activation free energy values Δ*G*‡(*T*) calculated using EVB/MD simulations for trypsins from (orange) warm-adapted (pig) and (blue) cold-adapted (arctic salmon) organisms, from Fig. 1 of (Isaksen et al., 2014). Solid lines are fits assuming temperature-independent Δ*H*‡ and Δ*S*‡. Dashed lines are fits to the cold-adapted results assuming **B** the same temperature-independent Δ*H*‡ as for the warm adapted variant and a temperature-dependent entropy Δ*S*‡(*T*) ; and **C** the same temperature-independent Δ*S*‡ as for the warm adapted variant and a temperature-dependent Δ*H* ‡(*T*). The fits in **A, B** and **C** are three of a manifold of possibilities that are consistent with the calculated Δ*G*‡(*T*).

An alternative approach to fitting the calculated Δ*G*‡(*T*) and experimental Δ*G*^*rxn*^ (*T*) is to relax the assumption that both entropy and enthalpy are temperature independent. This seems particularly appropriate for psychrophiles that show extensive surface mobility away from the active site and so are not as conformationally “locked down” as their meso- and thermophilic relatives. The constraint in this case is that Δ*H* ‡(*T*) and Δ*S*‡(*T*) at each temperature give the calculated Δ*G*‡(*T*). **Figure 5B** shows a fit to the calculated Δ*G*‡(*T*) for the cold-adapted variant assuming that Δ*H*‡ is constant and equal to that of the warm-adapted variant, and that Δ*S*‡(*T*) has a linear temperature dependence. The required entropy value and its temperature variation are not *a priori* unreasonable; for example, flexible surface residues in the cold-adapted version may adopt (bound) conformations in the transition state with fewer ordered waters, and as temperature increases the occupancy of those conformations may decrease. **Figure 5C** shows a fit to the calculated Δ*G*‡(*T*) for the cold-adapted variant assuming that Δ*H* ‡(*T*) varies linearly with temperature and that Δ*S*‡ is constant and equal to that of the warm-adapted variant. The enthalpy Δ*H* ‡(*T*) in this case is very close to that of the warm-adapted variant, and its change over the 35 °C range of the simulation results is only 1.1 kcal/mol, comparable to the energy of a single H bond. The fits in **Figs. 5 A, B** and **C** are just three of a manifold of possibilities that are consistent with the calculated Δ*G*‡(*T*). They suggest that the large difference in enthalpies between cold and warm adapted enzyme variants, especially those showing largely identical active sites, may be an artifact of assuming that Δ*H*‡ and Δ*S*‡ in the simulation (or Δ*H*^*rxn*^ and Δ*S*^*rxn*^ in the enzyme) are independent of temperature.

### 2.5 Thermodynamic constraints and Eyring fit parameters

Some interpretations of kinetics measurements assume that the enthalpy Δ*H* ^*rxn*^ (*T*) and entropy Δ*S*^*rxn*^ (*T*) governing the measured values are properties of a single transition state and so are subject to thermodynamic constraints (Arcus & Mulholland, 2020; Arcus, van der Kamp, Pudney, & Mulholland, 2020) (**SI Section S2**). Changes in enthalpy and entropy with temperature are connected to a heat capacity by

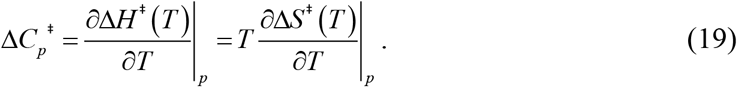

In general, the heat capacity 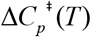 will be temperature dependent, and in the narrow absolute temperature range of biological relevance can be approximated as

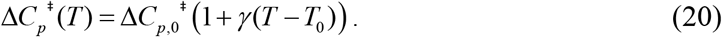

One interpretation (Arcus et al., 2020) assumes that 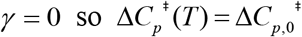. In that case, the Eyring relation becomes (Arcus & Mulholland, 2020)

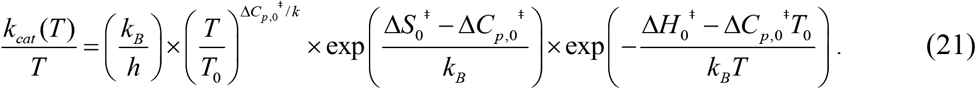

The term before the first exponential generates curvature in the corresponding Eyring plot. For a modest (e.g., biological) temperature range Δ*T* /*T*_0_ ≪ 1 around *T*_0_, Δ*S* ^‡^ (*T*) varies linearly with Δ*H* ^‡^ (*T*) (**SI Sec. S2**). In general, 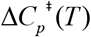 will be temperature dependent with *γ* ≠ 0 in Eq. 20, and Δ*H* ^‡^ (*T*), Δ*S* ^‡^ (*T*) and Δ*G*^‡^ (*T*) will vary nonlinearly with temperature (Walker et al., 2024).

A key question is whether quantities obtained by applying these nonlinear fits are physically meaningful. Enzymatic catalysis is in general a multi-step process. Each process can be characterized by its own thermodynamic parameters, each with its own temperature variations (Winzor & Jackson, 2006). For example, to describe a reaction with two steps after binding requires 2 sets of Δ*H* ^‡^ (*T*) and Δ*S* ^‡^ (*T*)< values, the temperature dependence of which may be described by one or more parameters, for a total of at least 6 parameters (Walker et al., 2024). While the heat capacity constraint Eq. 20 may apply to each of the underlying Δ*H* ^‡^ (*T*), Δ*S* ^‡^ (*T*) pairs, it may not apply to Δ*H* ^*rxn,app*^ (*T*) and Δ*S* ^*rxn,app*^ (*T*) values obtained by fitting experimental data with a simplified model such as Eq. 21.

## 3. Discussion

Enzymes have evolved to increase reaction rates at biological temperatures and to tailor the temperature dependence of those rates to achieve optimal behavior in a complex set of reactions as temperature varies. To do so, they have developed optimized active site geometries using flexible scaffolds and elements that undergo conformational changes to promote the reaction. A conformational change at high temperature – unfolding – causes large changes in activity, activation energy, and enthalpic and entropic barriers for the reaction. Changes in the conformational ensemble, in the active site geometry, and in distances and interaction strengths at lower temperatures throughout the biological temperature range should similarly produce temperature variation of the parameters (*A, E*_*a*_, Δ*H* and Δ*S*) describing the reaction. Temperature-dependent structural studies have provided evidence of such changes.

As shown here, linear Arrhenius/Eyring plots can be obtained even when the underlying parameters ((*A, E*_*a*_, Δ*H* and Δ*S*) vary with temperature. How does the usual assumption of *E* temperature-independent parameters affect interpretation of enzyme data? With modest variation of *E*_*a*_ (*T*), the apparent activitation energy 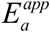 and prefactor *A*^*app*^ obtained from an Arrhenius fit may differ by a factor of ∼2 and by several orders of magnitude, respectively, from their underlying values. A temperature-dependent *E*_*a*_ (*T*) provides a generic, mechanism-agnostic explanation for anomalously large Arrhenius prefactors.

Similarly, an Eyring plot gives Δ*G*^*rxn*^ (*T*). Even when the plot appears linear, the values Δ*H* ^*rxn*^ (*T*) and Δ*S*^*rxn*^ (*T*) cannot be determined unless it is known that both are temperature independent. In the more likely case that one or both varies with temperature and so modulates the slope of Δ*G*^*rxn*^ (*T*), linear fit values Δ*H*^*rxn,app*^ and Δ*S*^*rxn,app*^ may be far from underlying values Δ*H*^*rxn*^ and Δ*S*^*rxn*^. When activation free energies Δ*G*‡(*T*) are calculated using, e.g., QM/MM simulations, fits assuming temperature-independent Δ*H*‡ and Δ*S*‡ may yield values far from underlying values and exhibit trends that do not, in fact, exist.

Consequently, expected small fractional variations of underlying thermodynamic /reaction parameters with temperature over the narrow biologically relevant range of absolute temperature will, in the absence of evidence from other measurements or simulations, make interpretation of parameters derived from Arrhenius /Eyring fits to enzyme kinetic data unreliable. Similar enzymes (e.g., mutants or isozymes) may have underlying free energy barriers Δ*G*^*rxn*^ (*T*) of comparable magnitude but yield very different Δ*H*^*rxn,app*^ and Δ*S*^*rxn,app*^ from fits to experimental data because of small differences in how their energy landscapes and conformational ensembles vary with temperature. Even qualitative ordering of, e.g., mutant enzymes based on their Arrhenius /Eyring slopes and/or intercepts may be problematic (Allen, Blum, Cunningham, Tu, & Hofmann, 1990; Lam, Yeung, Yu, Sze, & Wong, 2011; Lenaz, Sechi, Parenti-Castelli, Landi, & Bertoli, 1972; Lonhienne, Baise, Feller, Bouriotis, & Gerday, 2001; Nagel et al., 2011; Silverstein, 2012).

Moreover, the underlying *A(T), E*_*a*_ or Δ*H*, Δ*S* cannot be determined from Arrhenius/Eyring fits, except in the special case when both parameters are independent of temperature. In the general case, Arrhenius/Eyring fits will constrain their values and temperature variation but will not uniquely determine them.

### 3.1 Origins of enthalpy-entropy compensation in biological and non-biological systems

Enthalpy-entropy compensation is widely observed in fit parameters extracted from both kinetics and equilibrium measurements of enzyme systems and is also observed in Arrhenius/Eyring fit parameters for a wide variety of other biological and non-biological data. In the case of enzymes, simulations and theoretical arguments suggest that offsetting changes in enthalpy and entropy may be common. However, fit parameters from related enzymes often tightly adhere to a straight line with small residuals. Such precise apparent compensation is unlikely to be “real”, and points to a key role for non-mechanistic causes. Statistical uncertainty in measurements and fitting is one such cause (Boots & De Bokx, 1989; Cornish-Bowden, 2002, 2017; Exner, 2007; Krug et al., 1976a; Starikov & Nordén, 2007). In a two parameter linear fit, uncertainty in one parameter will cause “compensating” uncertainty in the other (Cornish-Bowden, 2018; Khrapunov, 2018). Algorithms have been proposed for ruling out statistical uncertainty as a cause of EEC in a given dataset (Griessen & Dam, 2021; Krug et al., 1976b, 1976a).

The present analysis suggests another general cause for apparent EEC, related to previously presented ideas (Cornish-Bowden, 2018; Khrapunov, 2018). Uncertainty here is not statistical as previously discussed but based on errors in the modelling required to extract thermodynamic parameters. Changes in thermodynamic potentials are linearly related to changes in thermodynamic variables, as in Gibbs relation Δ*G* = Δ*H* − *T* Δ*S*. In favorable cases, a given measurement may robustly determine one of the three quantities with minimal assumptions about mechanism. The Gibbs relation then constrains the other two quantities to lie upon a line. Determining a second quantity (and thus the third) from measurements often requires mechanistic assumptions and so is less robust. To the extent that otherwise similar enzyme/substrate systems exhibit larger and smaller deviations from the assumed behavior/model, the values of the second quantity will be dispersed relative to its underlying values, creating dispersion in calculated values of the third quantity. A plot of the second and third quantities will then show linear dispersion with changes in one associated with changes in the other.

For example, in activity measurements, *k*_*cat*_ (*T*) robustly determines Δ*G*^*rxn*^ (*T*) but only constrains Δ*H* ^*rxn*^ (*T*) and Δ*S*^*rxn*^ (*T*) to lie upon a straight line given by the Gibbs relation. An additional assumption — e.g., that Δ*H* ^*rxn*^ (*T*) and Δ*S*^*rxn*^ (*T*) are independent of temperature when Eyring plots appear linear — is required to obtain a unique solution. Deviations to varying extent in the underlying behavior from this assumption in otherwise similar enzymes will lead to deviations of fit parameters from underlying values, dispersed along a line, and apparent enthalpy-entropy compensation.

Similar considerations apply to equilibrium measurements of, e.g, binding and molecular recognition and their analysis using van ‘t Hoff plots. *K*_*eq*_ (*T*) robustly determines the standard free energy Δ*G*^0^ (*T*), but Δ*H* ^0^ (*T*) and Δ*S* ^0^ (*T*) are only constrained by the Gibbs equation. Linear van ‘t Hoff plots can occur even when the underlying Δ*H* ^0^ (*T*) and Δ*S* ^0^ (*T*) vary with temperature. For related systems with similar Δ*G*^0^ (*T*) but differences in their temperature variation, the apparent enthalpy and entropy values obtained from van’t Hoff plots will show correlated scatter.

Equilibrium measurements versus temperature have largely been supplanted by isothermal titration calorimetry (ITC). ITC measurements and fits are less susceptible to statistical errors, but EEC is still widely observed (Fox et al., 2017). The situation here is related to that for Arrhenius/Eyring and van’t Hoff analysis but is more complex. In ITC the heat released as a ligand is titrated is measured. Extracting the number of binding sites *n* per enzyme, enthalpy Δ*H*, and binding affinity *K*_*a*_ requires nonlinear fitting to specific models of binding (Bastos et al., 2023; Tellinghuisen, 2006). Δ*G* is calculated from the binding affinity *K*_*a*_, and Δ*S* calculated using the Gibbs relation. To the extent that the binding model assumptions fail to describe the actual binding for a given enzyme, the thermodynamic parameters deduced will deviate from their underlying values. These deviations will be correlated via the Gibbs relation, giving rise to apparent EEC. In addition, obtaining an analyzable titration curve within constraints of solubility and instrument sensitivity limits the range of measurable Δ*G* values, collapsing variability along that axis and leading to a rough linear correlation between Δ*H* and Δ*S* (Cooper, Johnson, Lakey, & Nollmann, 2001; Olsson, Ladbury, Pitt, & Williams, 2011). Binding in enzymes — which have complex, labile, and allosterically modulated active sites — may deviate substantially from models used in ITC fits, and the smooth titration curves obtained from ITC may not allow robust fitting to more sophisticated models.

Correlation /compensation between extracted Arrhenius parameters *A* and *E*_a_ has been observed in modeling many phenomena in non-biological systems over the last 70 years including chemical modification of organic compounds (Krug et al., 1976b, 1976a), thermal aging of polymers (Audouin & Verdu, 1991; Ernst, 1985), thermal decomposition of solids (Koga & Tanaka, 1991), and heterogeneous catalysis (Bond, 1981; Liu & Guo, 2001). These are all complex processes for which there is no reason to expect models involving a single temperature-independent activation energy to be adequate. In addition to experimental (statistical) errors, compensation has been shown to arise from deviations in the simple models used to fit and analyze the data from more physically plausible models, and from deviations in details of the latter models from the experimental system (Bond, 1981; Koga & Tanaka, 1991; Ernst, 1985; Audouin & Verdu, 1991; Liu & Guo, 2001).

In biological systems, Arrhenius /van’t Hoff parameter correlations and extremely large Arrhenius prefactors *A* have also been reported in protein denaturation and thermal degradation of cells and tissues (Qin et al., 2014; Wright, 2003). Physical mechanisms that may give large prefactors based on “loose” transition states have been proposed (Baklanov & Kiselev, 2023; Robinson, P. J. & Holbrook, K. A., 1972). However, reported Arrhenius prefactors of ∼10^129^ and 10^258^ for proteins and cells, respectively, co-occur with improbably high activation energies of ∼830 kcal/mol and 1830 kcal/mol, the former comparable to the total folding enthalpy of a ∼1000 residue protein. As with the non-biological phenomena cited above, thermal unfolding and especially cellular degradation are complex phenomena that are unlikely to be fully captured by simple two-state, single activation energy models in most cases. Deviations in behavior from those models may then account in part for observed parameter correlations.

## 4. Conclusion

The core problem, as has been previously pointed out (e.g., in (Winzor & Jackson, 2006)), is that parameters extracted from *k*_*cat*_ (*T*) data (or *K*_*eq*_ (*T*) data or ITC data) using any model that does not closely represent the underlying enzymatic mechanism are just parameters. Assumptions used in analyzing small molecule systems may not account for the complexity and temperature variation of enzyme action. Even small deviations from assumed behavior can have a large impact on extracted parameters and even qualitative trends in parameters extracted from related enzymes may be suspect. Models that more accurately represent the underlying mechanism may have so many parameters that fitting of measurements obtained using a single technique will be insufficient to determine them. All of this points to a critical role for use of complementary methods including QM/MM simulations and multi-temperature high-resolution static and time-resolved studies of enzyme structure, ligand interactions, and function to guide modeling and interpretation.

## Acknowledgments

None.

## Author contributions

Conceptualization: MJM, RET

Methodology: RET

Investigation: RET

Analysis: RET

Funding acquisition: RET

Project administration: RET

Supervision: RET

Writing – original draft: RET

Writing – review & editing: MJM, RET,

## References

Allen, B., Blum, M., Cunningham, A., Tu, G. C., & Hofmann, T. (1990). A ligand-induced, temperature-dependent conformational change in penicillopepsin. Evidence from nonlinear Arrhenius plots and from circular dichroism studies. Journal of Biological Chemistry, 265(9), 5060–5065. doi: 10.1016/s0021-9258(19)34084-0

Åqvist, J., & Brandsdal, B. O. (2025). Computer Simulations of the Temperature Dependence of Enzyme Reactions. Journal of Chemical Theory and Computation, 21(3), 1017–1028. doi: 10.1021/acs.jctc.4c01733

Åqvist, J., Isaksen, G. V., & Brandsdal, B. O. (2017). Computation of enzyme cold adaptation. Nature Reviews Chemistry, 1. doi: 10.1038/s41570-017-0051

Arcus, V. L., & Mulholland, A. J. (2020). Temperature, Dynamics, and Enzyme-Catalyzed Reaction Rates. Annual Review of Biophysics, 49, 163–180. doi: 10.1146/annurev-biophys-121219

Arcus, V. L., van der Kamp, M. W., Pudney, C. R., & Mulholland, A. J. (2020). Enzyme evolution and the temperature dependence of enzyme catalysis. Current Opinion in Structural Biology, 65, 96–101. doi: 10.1016/j.sbi.2020.06.001

Arrhenius, S. (1889). Über die Dissociationswärme und den Einfluss der Temperatur auf den Dissociationsgrad der Elektrolyte. Zeitschrift Für Physikalische Chemie, 4(1), 96–116. doi: 10.1515/zpch-1889-0408

Audouin, L., & Verdu, J. (1991). Comments on the Electrotechnical Ageing Compensation Effect. Polymer Degradation and Stability, 31, 335–346. doi: 10.1016/0141-3910(91)90041-O

Baklanov, A. V., & Kiselev, V. G. (2023). The Nature of the Enthalpy–Entropy Compensation and “Exotic” Arrhenius Parameters in the Denaturation Kinetics of Proteins. International Journal of Molecular Sciences, 24(13), 10630. doi: 10.3390/ijms241310630

Barrie, P. J. (2012). The mathematical origins of the kinetic compensation effect: 2. The effect of systematic errors. Physical Chemistry Chemical Physics, 14(1), 327–336. doi: 10.1039/c1cp22667c

Bastos, M., Abian, O., Johnson, C. M., Ferreira-da-Silva, F., Vega, S., Jimenez-Alesanco, A., … Velazquez-Campoy, A. (2023). Isothermal titration calorimetry. Nature Reviews Methods Primers, 3(1), 17. doi: 10.1038/s43586-023-00199-x

Bond, G. C. (1981). The significance of the compensation effect and the definition of active centres in metal catalysts. Zeitschrift Fur Physikalische Chemie Neue Folge Bd., 144, 21–31. doi: 10.1524/zpch.1985.144.144.021

Boots, H. M. J., & De Bokx, P. K. (1989). Theory of Enthalpy-Entropy Compensation. J. Phys. Chem, 93, 8240– 8243. doi: 10.1021/j100362a018

Chodera, J. D., & Mobley, D. L. (2013). Entropy-Enthalpy Compensation: Role and Ramifications in Biomolecular Ligand Recognition and Design. Annual Review of Biophysics, 42(1), 121–142. doi: 10.1146/annurev-biophys-083012-130318

Cooper, A., Johnson, C. M., Lakey, J. H., & Nollmann, M. (2001). Heat does not come in different colours: Entropy-enthalpy compensation, free energy windows, quantum confinement, pressure perturbation calorimetry, solvation and the multiple causes of heat capacity effects in biomolecular interactions. Biophysical Chemistry, 93, 215–230. doi: 10.1016/S0301-4622(01)00222-8

Cornish-Bowden, A. (2002). Enthalpy-entropy compensation: A phantom phenomenon. J. Biosci., 27(2), 121–126. doi: 10.1007/BF02703768

Cornish-Bowden, A. (2017). Enthalpy–entropy compensation and the isokinetic temperature in enzyme catalysis. Journal of Biosciences, 42(4), 665–670. doi: 10.1007/s12038-017-9719-0

Cornish-Bowden, A. (2018). Entropy-Enthalpy Compensation. In European Biophysical Societies, G. Roberts, & A. Watts (Eds.), Encyclopedia of Biophysics (pp. 1–6). Berlin, Heidelberg: Springer Berlin Heidelberg. doi: 10.1007/978-3-642-35943-9_10072-1

Ernst, W. R. (1985). False compensation effect in studies of kinetics of cure by differential scanning calorimetry. Thermochimica Acta, 91, 127–133. doi: 10.1016/0040-6031(85)85208-4

Evans, M. G., & Polanyi, M. (1935). Some Applications of the Transition State Method to the Calculation of Reaction Velocities, Especially in Solution. Transactions of the Faraday Society, 31, 875–894. doi: 10.1039/TF9353100875

Exner, O. (1973). The Enthalpy-Entropy Relationship. In Progress in Physical Organic Chemistry (Vol. 10, pp. 411–482). Wiley. doi: 10.1002/9780470171899.ch6

Exner, O. (2007). The Enthalpy-Entropy Relationship. Progress in Physical Organic Chemistry, 10, 411–482. doi: 10.1002/9780470171899.ch6

Eyring, H. (1935a). The Activated Complex in Chemical Reactions. The Journal of Chemical Physics, 3(2), 63–71. doi: 10.1063/1.1749604

Eyring, H. (1935b). The Activated Complex in Chemical Reactions. The Journal of Chemical Physics, 3(2), 107– 115. doi: 10.1063/1.1749604

Eyring, Henry. (1935c). The Activated Complex and the Absolute Rate of Chemical Reactions. Chemical Reviews, 17(1), 65–77. doi: 10.1021/cr60056a006

Feller, G., & Gerday, C. (2003). Psychrophilic enzymes: Hot topics in cold adaptation. Nature Reviews Microbiology, 1, 200–208. doi: 10.1038/nrmicro773

Fields, P. A., & Somero, G. N. (1998). Hot spots in cold adaptation: Localized increases in conformational flexibility in lactate dehydrogenase A4 orthologs of Antarctic notothenioid fishes. Proceedings of the National Academy of Sciences, 95(19), 11476–11481. doi: 10.1073/pnas.95.19.11476

Fox, J. M., Zhao, M., Fink, M. J., Kang, K., & Whitesides, G. M. (2017). The Molecular Origin of Enthalpy/Entropy Compensation in Biomolecular Recognition. Annual Review of Biophysics, 47, 1–28. doi: 10.26434/chemrxiv.5649775.v1

Garcia-Viloca, M., Gao, J., Karplus, M., & Truhlar, D. G. (2004). How Enzymes Work: Analysis by Modern Rate Theory and Computer Simulations. Science, 303(5655), 186–195. doi: 10.1126/science.1088172

Griessen, R., & Dam, B. (2021). Simple Accurate Verification of Enthalpy-Entropy Compensation and Isoequilibrium Relationship. ChemPhysChem, 22(17), 1774–1784. doi: 10.1002/cphc.202100431

Herschlag, D., & Pinney, M. M. (2018). Hydrogen Bonds: Simple after All? Biochemistry, 57(24), 3338–3352. doi: 10.1021/acs.biochem.8b00217

Hynes, J. T., Laage, D., Tuñón, I., & Moliner, V. (2016). Chapter 3. A Transition State Theory Perspective for Enzymatic Reactions: Fundamentals and Applications. In I. Tunon & V. Moliner (Eds.), Theoretical and Computational Chemistry Series (pp. 54–88). Cambridge: Royal Society of Chemistry. doi: 10.1039/9781782626831-00054

Isaksen, G. V., Åqvist, J., & Brandsdal, B. O. (2014). Protein Surface Softness Is the Origin of Enzyme Cold-Adaptation of Trypsin. PLoS Computational Biology, 10(8), e1003813. doi: 10.1371/journal.pcbi.1003813

Isaksen, G. V., Åqvist, J., & Brandsdal, B. O. (2016). Enzyme surface rigidity tunes the temperature dependence of catalytic rates. PNAS, 113(28), 7822–7827. doi: 10.1073/pnas.1605237113

Kazemi, M., & Åqvist, J. (2015). Chemical reaction mechanisms in solution from brute force computational Arrhenius plots. Nature Communications, 6(1), 7293. doi: 10.1038/ncomms8293

Keedy, D. A., Hill, Z. B., Biel, J. T., Kang, E., Rettenmaier, T. J., Brandão-Neto, J., … Fraser, J. S. (2018). An expanded allosteric network in PTP1B by multitemperature crystallography, fragment screening, and covalent tethering. eLife, 7, e36307. doi: 10.7554/eLife.36307

Keedy, D. A., Kenner, L. R., Warkentin, M., Woldeyes, R. A., Hopkins, J. B., Thompson, M. C., … Fraser, J. S. (2015). Mapping the conformational landscape of a dynamic enzyme by multitemperature and XFEL crystallography. eLife, 4, 07574. doi: 10.7554/eLife.07574

Khrapunov, S. (2018). The Enthalpy-Entropy Compensation Phenomenon. Limitations for the Use of Some Basic Thermodynamic Equations. Current Protein & Peptide Science, 19(11), 1088–1091. doi: 10.2174/1389203719666180521092615

Koga, N., & Tanaka, H. (1991). A kinetic compensation effect established for the thermal decomposition of a solid. Journal of Thermal Analysis, 37(2), 347–363. doi: 10.1007/BF02055937

Kohen, Amnon, Cannio, Raffaele, Bartolucci, Simonetta, & Klinman Judith P. (1999). Enzyme dynamics and hydrogen tunneling in a thermophilic alchohol dehydrogenase. Nature, 399, 496–499. doi: doi.org/10.1524/zpch.1985.144.144.021

Krug, R. R., Hunter, W. G., & Grieger, R. A. (1976a). Enthalpy-entropy compensation. 1. Some fundamental statistical problems associated with the analysis of van’t Hoff and Arrhenius data. The Journal of Physical Chemistry, 80(21), 2335–2341. doi: 10.1021/j100562a006

Krug, R. R., Hunter, W. G., & Grieger, R. A. (1976b). Statistical interpretation of enthalpy–entropy compensation. Nature, 261(5561), 566–567. doi: 10.1038/261566a0

Lam, S. Y., Yeung, R. C. Y., Yu, T. H., Sze, K. H., & Wong, K. B. (2011). A rigidifying salt-bridge favors the activity of thermophilic enzyme at high temperatures at the expense of low-temperature activity. PLoS Biology, 9(3). doi: 10.1371/journal.pbio.1001027

Lenaz, G., Sechi, A. M., Parenti-Castelli, G., Landi, L., & Bertoli, E. (1972). Activation Energies of Different Mitochondrial Enzymes: Breaks in Arrhenius Plots of Membrane-Bound Enzymes Occur at Different Temperatures. Biochemical and Biophysical Research Communications, 49(2), 536–542. doi: 10.1016/0006-291X(72)90444-5

Liu, L., & Guo, Q.-X. (2001). Isokinetic Relationship, Isoequilibrium Relationship, and Enthalpy−Entropy Compensation. Chemical Reviews, 101(3), 673–696. doi: 10.1021/cr990416z

Lonhienne, T., Baise, E., Feller, G., Bouriotis, V., & Gerday, C. (2001). Enzyme activity determination on macromolecular substrates by isothermal titration calorimetry: Application to mesophilic and psychrophilic chitinases. Biochimica et Biophysica Acta, 1545, 349–456. doi: 10.1016/S0167-4838(00)00296-X

Machado, T. F. G., Gloster, T. M., & Da Silva, R. G. (2018). Linear Eyring Plots Conceal a Change in the Rate-Limiting Step in an Enzyme Reaction. Biochemistry, 57(49), 6757–6761. doi: 10.1021/acs.biochem.8b01099

McLeod, M. J., Barwell, S. A. E., Holyoak, T., & Thorne, R. E. (2025). A structural perspective on the temperature dependent activity of enzymes. Structure, 33(5), 924-934.e2. doi: 10.1016/j.str.2025.02.013

Nagel, Z. D., Dong, M., Bahnson, B. J., & Klinman, J. P. (2011). Impaired protein conformational landscapes as revealed in anomalous Arrhenius prefactors. PNAS, 108(26), 10520–10525. doi: 10.1073/pnas.1104989108

Olsson, T. S. G., Ladbury, J. E., Pitt, W. R., & Williams, M. A. (2011). Extent of enthalpy–entropy compensation in protein–ligand interactions. Protein Science, 20(9), 1607–1618. doi: 10.1002/pro.692

Pedersen, H. L., Willassen, N. P., & Leiros, I. (2009). The first structure of a cold-adapted superoxide dismutase (SOD): Biochemical and structural characterization of iron SOD from Aliivibrio salmonicida. Acta Crystallographica Section F Structural Biology and Crystallization Communications, 65(2), 84–92. doi: 10.1107/S1744309109001110

Peters, B. (2017). Reaction rate theory and rare events. Amsterdam ; Cambrige, MA: Elsevier.

Purich, D. L. (2010). Enzyme Kinetics. Elsevier. doi: 10.1016/B978-0-12-380924-7.10003-1

Qin, Z., Balasubramanian, S. K., Wolkers, W. F., Pearce, J. A., & Bischof, J. C. (2014). Correlated Parameter Fit of Arrhenius Model for Thermal Denaturation of Proteins and Cells. Annals of Biomedical Engineering, 42(12), 2392–2404. doi: 10.1007/s10439-014-1100-y

Robinson, P. J. & Holbrook, K. A. (1972). Unimolecular Reactions. London: Wiley.

Siddiqui, K. S., & Cavicchioli, R. (2006). Cold-Adapted Enzymes. Annual Review of Biochemistry, 75(1), 403–433. doi: 10.1146/annurev.biochem.75.103004.142723

Silverstein, T. P. (2012). Falling enzyme activity as temperature rises: Negative activation energy or denaturation? Journal of Chemical Education, 89(9), 1097–1099. doi: 10.1021/ed200497r

Smalås, A. O., Heimstad, E. S., Hordvik, A., Willassen, N. P., & Male, R. (1994). Cold adaption of enzymes: Structural comparison between salmon and bovine trypsins. Proteins: Structure, Function, and Bioinformatics, 20(2), 149–166. doi: 10.1002/prot.340200205

Starikov, E. B., & Nordén, B. (2007). Enthalpy-entropy compensation: A phantom or something useful? Journal of Physical Chemistry B, 111(51), 14431–14435. doi: 10.1021/jp075784i

Tellinghuisen, J. (2006). Van’t Hoff analysis of K°(T): How good…or bad? Biophysical Chemistry, 120(2), 114– 120. doi: 10.1016/j.bpc.2005.10.012

Truhlar, D. G. (2015). Transition state theory for enzyme kinetics. Archives of Biochemistry and Biophysics, 582, 10–17. doi: 10.1016/j.abb.2015.05.004

Tsigos, I., Mavromatis, K., Tzanodaskalaki, M., Pozidis, C., Kokkinidis, M., & Bouriotis, V. (2001). Engineering the properties of a cold active enzyme through rational redesign of the active site. European Journal of Biochemistry, 268(19), 5074–5080. doi: 10.1046/j.0014-2956.2001.02432.x

Walker, E. J., Hamill, C. J., Crean, R., Connolly, M. S., Warrender, A. K., Kraakman, K. L., … Arcus, V. L. (2023). Cooperative conformational transitions and the temperature dependence of enzyme catalysis [Preprint]. Biochemistry. doi: 10.1101/2023.07.06.548038

Walker, E. J., Hamill, C. J., Crean, R., Connolly, M. S., Warrender, A. K., Kraakman, K. L., … Arcus, V. L. (2024). Cooperative Conformational Transitions Underpin the Activation Heat Capacity in the Temperature Dependence of Enzyme Catalysis. ACS Catalysis, 14(7), 4379–4394. doi: 10.1021/acscatal.3c05584

Winzor, D. J., & Jackson, C. M. (2006). Interpretation of the temperature dependence of equilibrium and rate constants. Journal of Molecular Recognition, 19(5), 389–407. doi: 10.1002/jmr.799

Wright, N. T. (2003). On a Relationship Between the Arrhenius Parameters from Thermal Damage Studies. Journal of Biomechanical Engineering, 125(2), 300–304. doi: 10.1115/1.1553974

