## Supplementary information for "Activation parameters, enthalpy-entropy compensation and the temperature-dependent activity of enzymes"

**This PDF file includes:**

Secs. S1 and S2
Figs. S1-4
Tables S1-5

### S1. Interpretation of hydride transfer measurements

Kinetics measurements of hydride transfer in WT and mutant thermophilic alcohol dehydrogenases indicated a break in Arrhenius slope at ~30 °C (Nagel et al., 2011). Above this temperature, all variants showed similar slopes and intercepts, and fits yielded physically reasonable Arrhenius prefactors. Below this temperature, slopes varied substantially between variants (Fig. 2 in Nagel *et al.*) and fits yielded unreasonable prefactors as large as $10^{25}$ $s^{-1}$. **Tables S2** gives the parameters $\Delta H^{rxn,app}$, $T\Delta S^{rxn,app}$, and $\log(A^{app})$ determined from linear fits, from Tables 2 and 3 in Nagel *et al.*

These data were accounted for by a model where the enzyme interconverts between competent and incompetent conformers separated by a free energy difference $\Delta G_c^0$ below the break, and where only competent conformers are populated above the break (due to, e.g., a conformational change that raised the energy of incompetent states relative to competent states.) Below 30 °C, the fraction of enzymes in the competent conformation was estimated, using a two-state model, as

$$f(T) = \frac{e^{-\Delta G_c/k_B T}}{1+e^{-\Delta G_c/k_B T}} \approx e^{-\Delta G_c/k_B T} \quad \text{(S1)}$$

where $\Delta G_c = \Delta H_c - T\Delta S_c$ is the free-energy difference between competent and incompetent conformations. The observed $k_{cat}(T)$ was then given by

$$k_{cat}(T) = e^{-\Delta G_c/k_B T} \times \left( \frac{k_B T}{h} e^{\Delta S^{rxn}/k_B} \right) e^{-\Delta H^{rxn}/k_B T}, \quad \text{(S2)}$$

where $\Delta H^{rxn}$ and $\Delta S^{rxn}$ are values for the competent conformation. **Table S3** gives the parameters $\Delta H_c$ and $T\Delta S_c$ obtained by applying this fit to data at temperatures below 30 °C, from Nagel *et al.*, Table 4. While $\Delta H_c$ is large – between 7 and 22 kcal/mol, of the same order as $\Delta H^{rxn,app}$ in **Table S2**, it is almost perfectly compensated by $T\Delta S_c$ so that $\Delta G_c \sim 0.1$ kcal/mol for all variants.

**S2. Thermodynamic constraints and Eyring fit parameters**

Kinetics measurements can be analyzed by assuming that the enthalpy and entropy governing the measured values are properties of a single (transition) state and so are subject to thermodynamic constraints (Arcus & Mulholland, 2020; Arcus et al., 2020). Changes in enthalpy and entropy with temperature are connected to a heat capacity by

$$\Delta C_p^{\ddagger} = \left.\frac{\partial \Delta H^{\ddagger}(T)}{\partial T}\right|_p = T\left.\frac{\partial \Delta S^{\ddagger}(T)}{\partial T}\right|_p . \tag{S3}$$

In general, the heat capacity $\Delta C_p^{\ddagger}(T)$ will be temperature dependent. In the narrow absolute temperature range of biological relevance, the first two terms of a Taylor series approximation give

$$\Delta C_p^{\ddagger}(T) = \Delta C_{p,0}^{\ddagger}\left(1+\gamma(T-T_0)\right). \tag{S4}$$

Macromolecular rate theory (MMRT) (Arcus et al., 2020) in its original form assumes that $\gamma = 0$ so $\Delta C_p^{\ddagger}(T) = \Delta C_{p,0}^{\ddagger}$ is temperature independent. In that case,

$$\Delta H^{\ddagger}(T) = \Delta H_0^{\ddagger} + \Delta C_{p,0}^{\ddagger}(T-T_0), \tag{S5}$$

$$\Delta S^{\ddagger}(T) = \Delta S_0^{\ddagger} + \Delta C_{p,0}^{\ddagger}\ln(T/T_0), \tag{S6}$$

and the Eyring relation becomes

$$\frac{k_{cat}(T)}{T} = \left(\frac{k_B}{h}\right)\times\exp\left(\frac{\Delta S_0^{\ddagger} - \Delta C_{p,0}^{\ddagger}}{k_B}\right)\times\left(\frac{T}{T_0}\right)^{\Delta C_{p,0}^{\ddagger}/k}\times\exp\left(-\frac{\Delta H_0^{\ddagger} - \Delta C_{p,0}^{\ddagger}T_0}{k_B T}\right). \tag{S7}$$

The term before the second exponential generates curvature in the resulting Eyring plot. Noting that for $\Delta T/T_0 = (T-T_0)/T_0 \ll 1$ (which is often the case in kinetics measurements), $\ln(T/T_0) = \ln\left(1+\Delta T/T_0\right) \approx \Delta T/T_0$, and the relation between $\Delta H^{\ddagger}(T)$ and $\Delta S^{\ddagger}(T)$ is then

$$\Delta S^{\ddagger}(T) \approx \left(\Delta S_0^{\ddagger} - \frac{\Delta H_0^{\ddagger}}{T_0}\right) + \frac{\Delta H^{\ddagger}(T)}{T_0}. \tag{S8}$$

The first term on the right is a constant. Thus, if $\Delta H^{\ddagger}(T)$ and $\Delta S^{\ddagger}(T)$ are connected by a constant heat capacity, $\Delta S^{\ddagger}(T)$ varies approximately linearly with $\Delta H^{\ddagger}(T)$. For a series of enzymes with different constant heat capacities $\Delta C_{p,0}{}^{\ddagger}$ and similar $\Delta G_0{}^{\ddagger} = \Delta H_0{}^{\ddagger} - T_0 \Delta S_0{}^{\ddagger}$ values, there will be enthalpy-entropy compensation (over a temperature range where the approximation above holds) with a compensation temperature comparable to what has been experimentally observed, even though the corresponding Eying plot will be nonlinear. The values of $\Delta C_{p,0}{}^{\ddagger}$ required to reproduce experimental curvatures generate large variations in $\Delta H^{\ddagger}(T)$ and $\Delta S^{\ddagger}(T)$; a fit to the data in Fig. 3 of (McLeod et al., 2025) requires a factor of ~4 variation in $\Delta H^{\ddagger}(T)$ and a factor of ~5 variation in $\Delta S^{\ddagger}(T)$ over the experimental temperature range.

In general, $\Delta C_p{}^{\ddagger}(T)$ will be temperature dependent. Assuming $\gamma \neq 0$ in Eq. S4, Eqs. S5 and S6 become

$$\Delta H^{\ddagger}(T) = \Delta H_0{}^{\ddagger} + \Delta C_{p,0}{}^{\ddagger}(T - T_0) + \frac{\gamma}{2} \Delta C_{p,0}{}^{\ddagger}(T - T_0)^2 \qquad \text{(S9)}$$

and

$$\Delta S^{\ddagger}(T) = \Delta S_0{}^{\ddagger} + \Delta C_{p,0}{}^{\ddagger}(1 - \gamma T_0)\ln(T / T_0) + \gamma \Delta C_{p,0}{}^{\ddagger}(T - T_0). \qquad \text{(S10)}$$

The additional terms add additional nonlinearity that can improve fits to $k_{cat}(T)$ data. This approximation corresponds to a recently updated version of MMRT, MMRT-1L (Walker et al., 2023).

### SI Figures

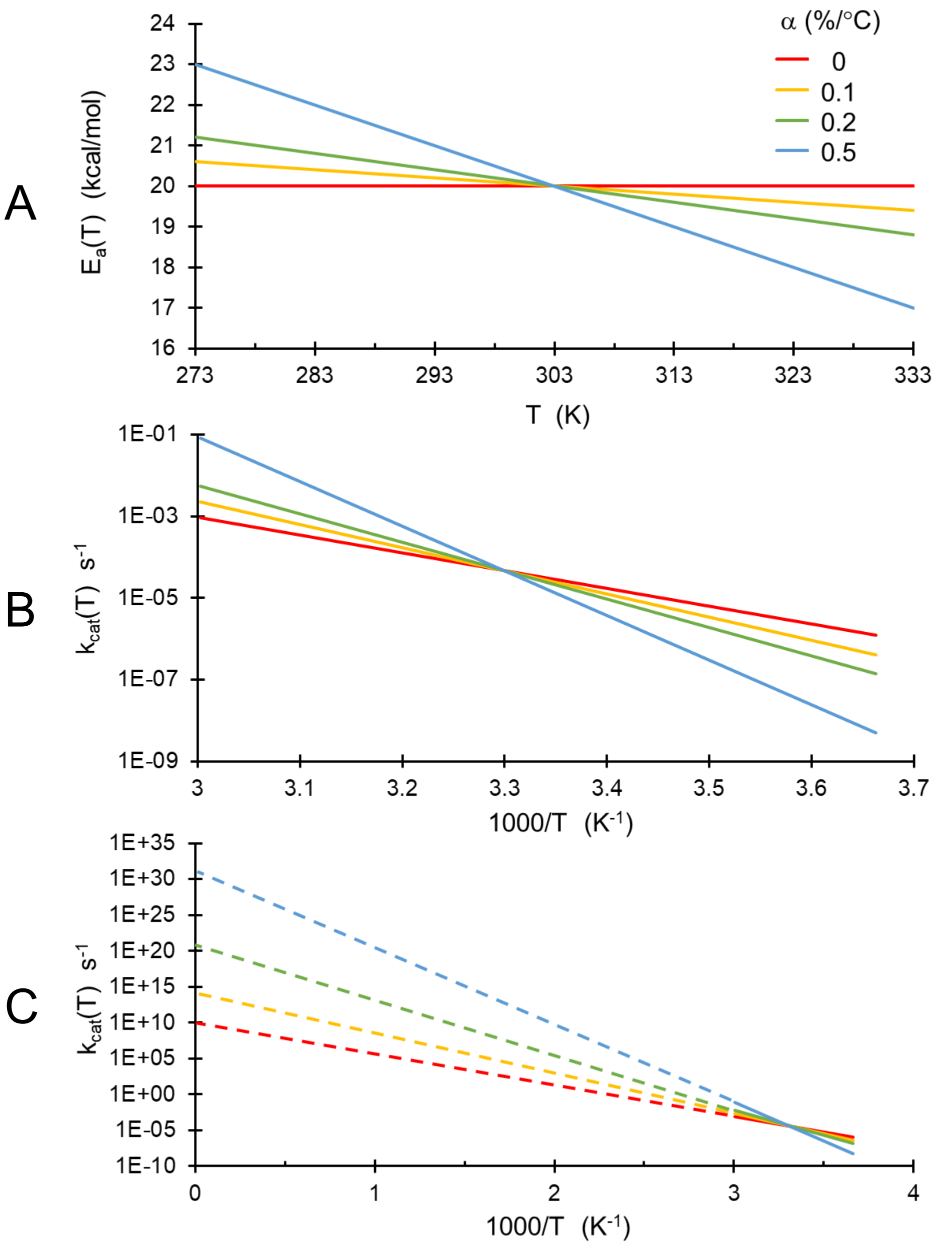


**Figure S1.** Effect of variations of activation energy $E_a(T)$ with temperature on $k_{cat}(T)$. $E_a(T=T_0)=20$ kcal/mol and $T_0=303$ K in all cases. **A** $E_a(T)$ vs $T$ for the first four values of the temperature coefficient $\alpha$ in Table S1. **B** Corresponding $k_{cat}(T)$ vs $1/T$ over a biological temperature range, with $A=10^{10}$ $s^{-1}$ in all cases. **C** Extrapolation of $k_{cat}(T)$ to $1/T=0$ to obtain the apparent Arrhenius prefactor $A^{app}$. Temperature variation of $E_a(T)$ of 12% over the biological temperature range increases the apparent prefactor by a factor of ~$10^{11}$.

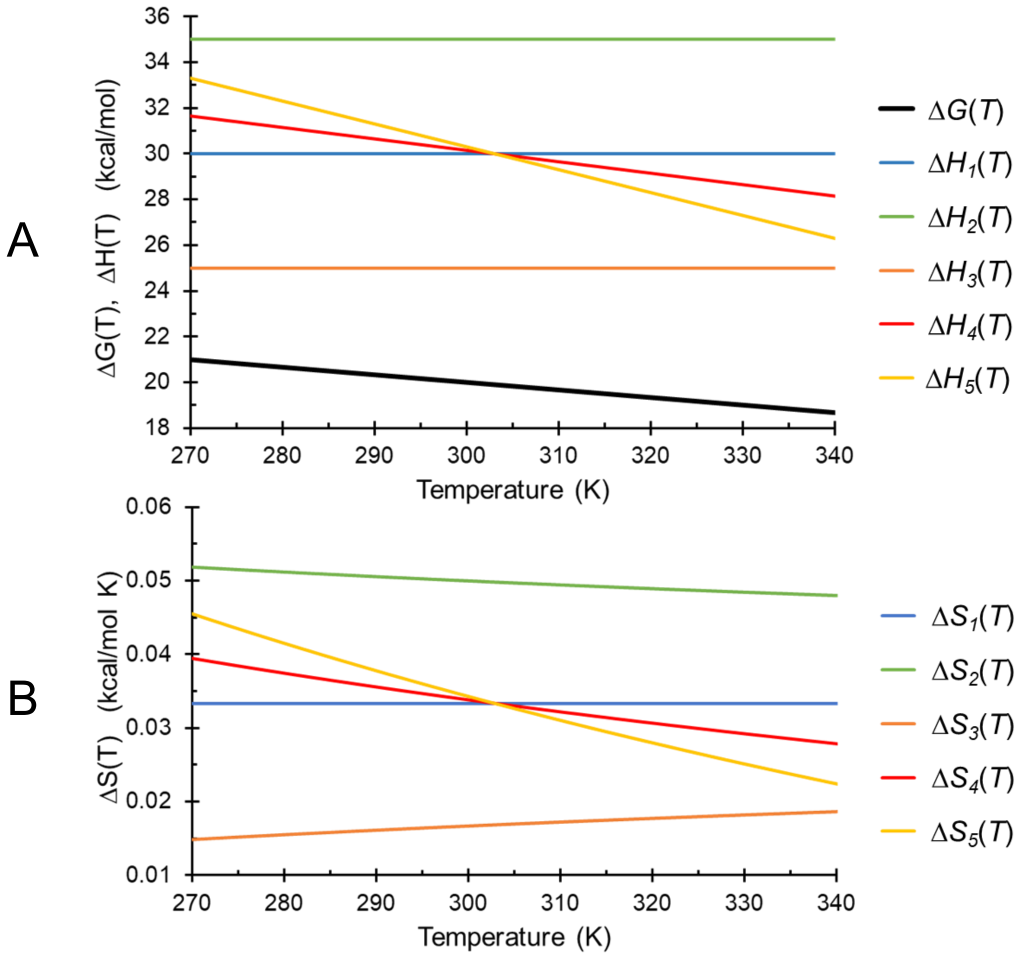


**Figure S2.** In the Eyring or van ‘t Hoff models, measurements of a kinetic constant $k_{cat}(T)$ or equilibrum constant $K_{eq}(T)$ versus $T$ directly give a temperature dependent free energy $\Delta G(T)$, related by the Gibbs relation $\Delta G(T) = \Delta H(T) - T\Delta S(T)$. If $\Delta H$ and $\Delta S$ are both independent of temperature and $\Delta G(T)$ varies linearly with temperature, then there is a unique solution to the Gibbs relation for $\Delta H$ and $\Delta S$. In general, the temperature dependence of $\Delta H$ and $\Delta S$ is unknown, and so their values cannot be determined. The figure shows a free energy $\Delta G(T)$ and examples of $\Delta H(T)$, $\Delta S(T)$ pairs that are consistent with the given $\Delta G(T)$. The 5 cases shown correspond to (1) constant $\Delta H$ and $\Delta S$; (2) and (3) $\Delta H$ constant, $\Delta S$ linearly varying; (4) and (5) $\Delta H$ and $\Delta S$ linearly varying. Nonlinear variations of $\Delta H$ and $\Delta S$ with temperature can also be consistent with a linear $\Delta G(T)$.

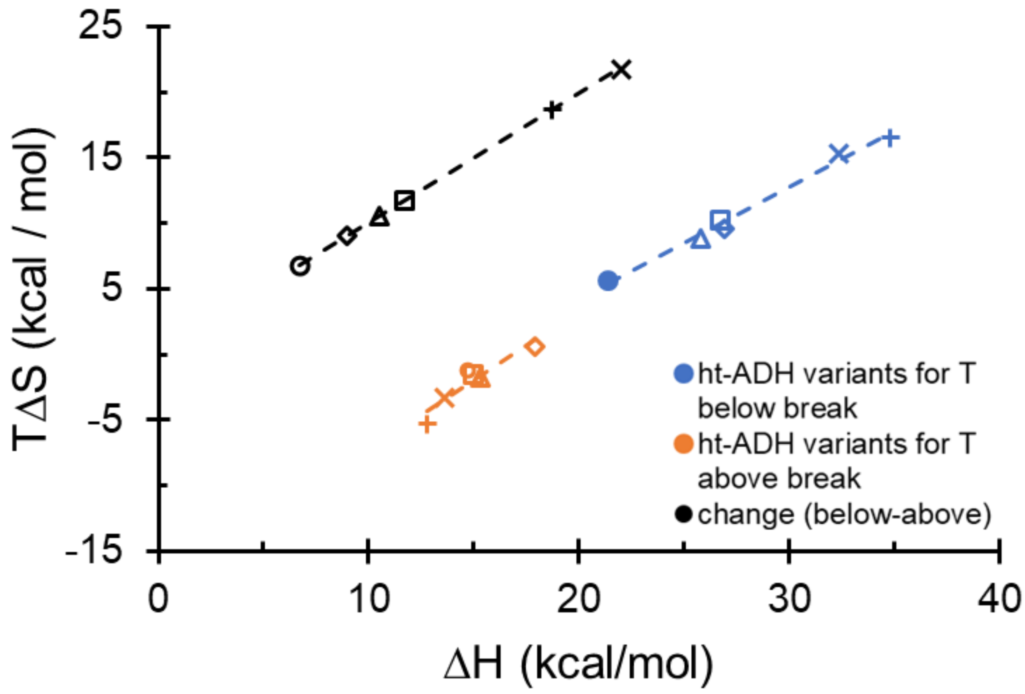


**Figure S3.** Parameters for Eyring fits to data above (orange) and below (blue) the break for six ht-ADH variants, from (Nagel et al., 2011). Linear fits give $R^2$=0.87 above the break and $R^2$=0.987 below the break. The changes in $\Delta H$ and $T\Delta S$ at the break (black) are nearly equal and opposite for all variants, giving nearly perfect compensation; the linear fit has $R^2$=0.9999.

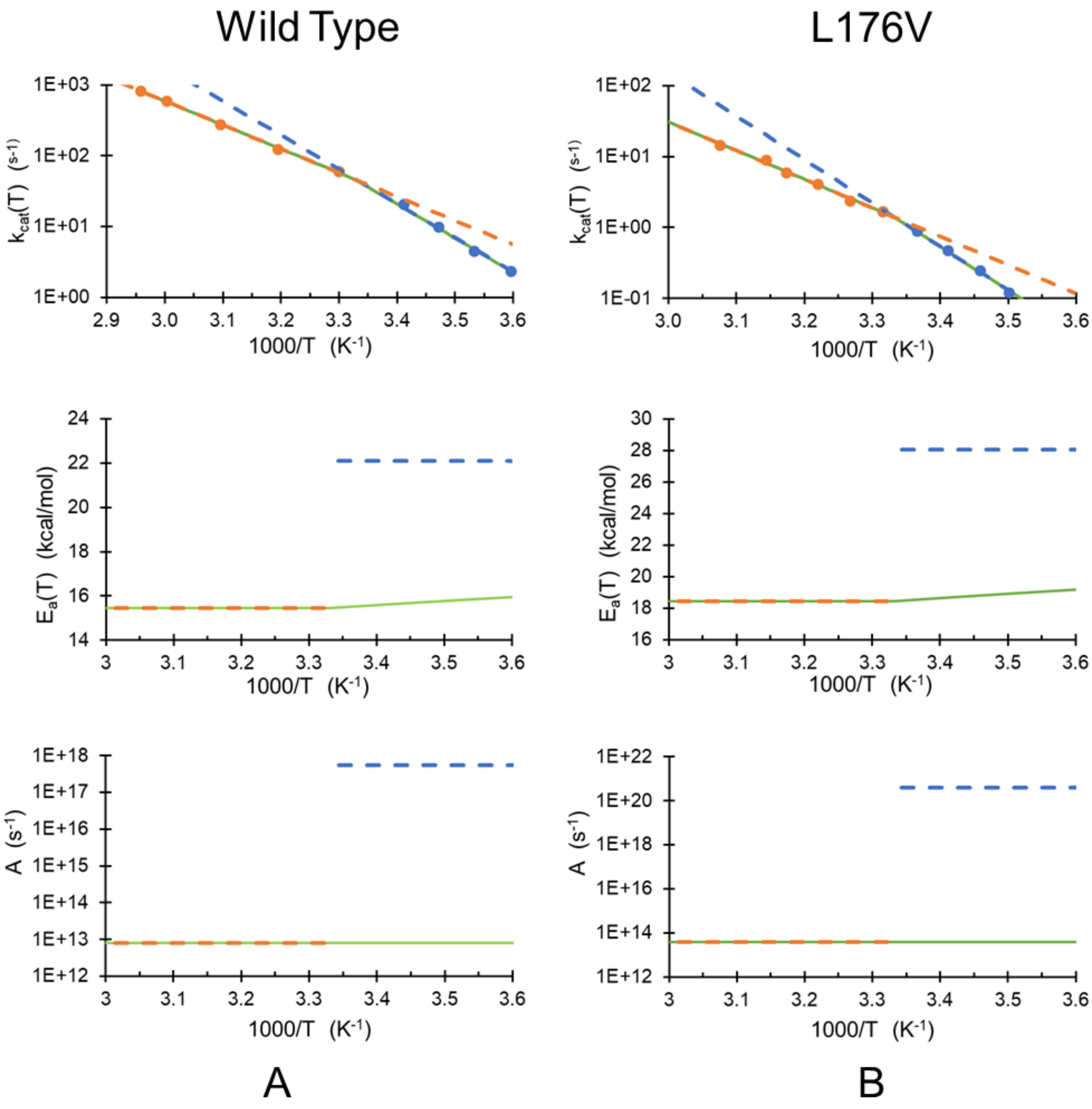

Wild Type
L176V
1E+03
1E+02
1E+01
1E+00
$k_{cat}(T)$ ($s^{-1}$)
2.9
3.0
3.1
3.2
3.3
3.4
3.5
3.6
1000/T ($K^{-1}$)
1E+02
1E+01
1E+00
1E-01
24
22
20
18
16
14
$E_a(T)$ (kcal/mol)
30
28
26
24
22
20
18
16
1E+18
1E+17
1E+16
1E+15
1E+14
1E+13
1E+12
A ($s^{-1}$)
1E+22
1E+20
1E+18
1E+16
1E+14
A
B


Fig. S4

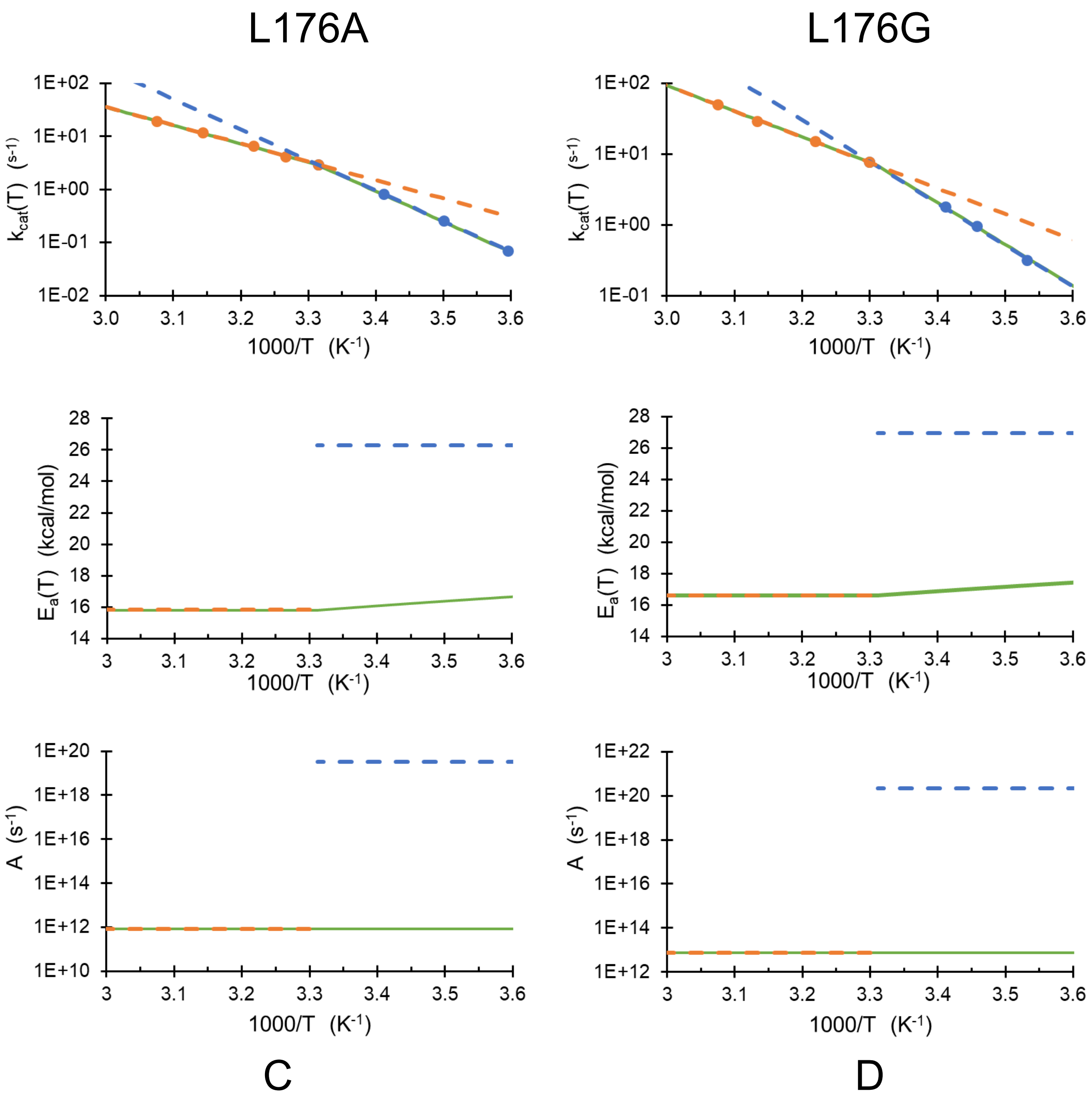


Fig. S4 (continued)

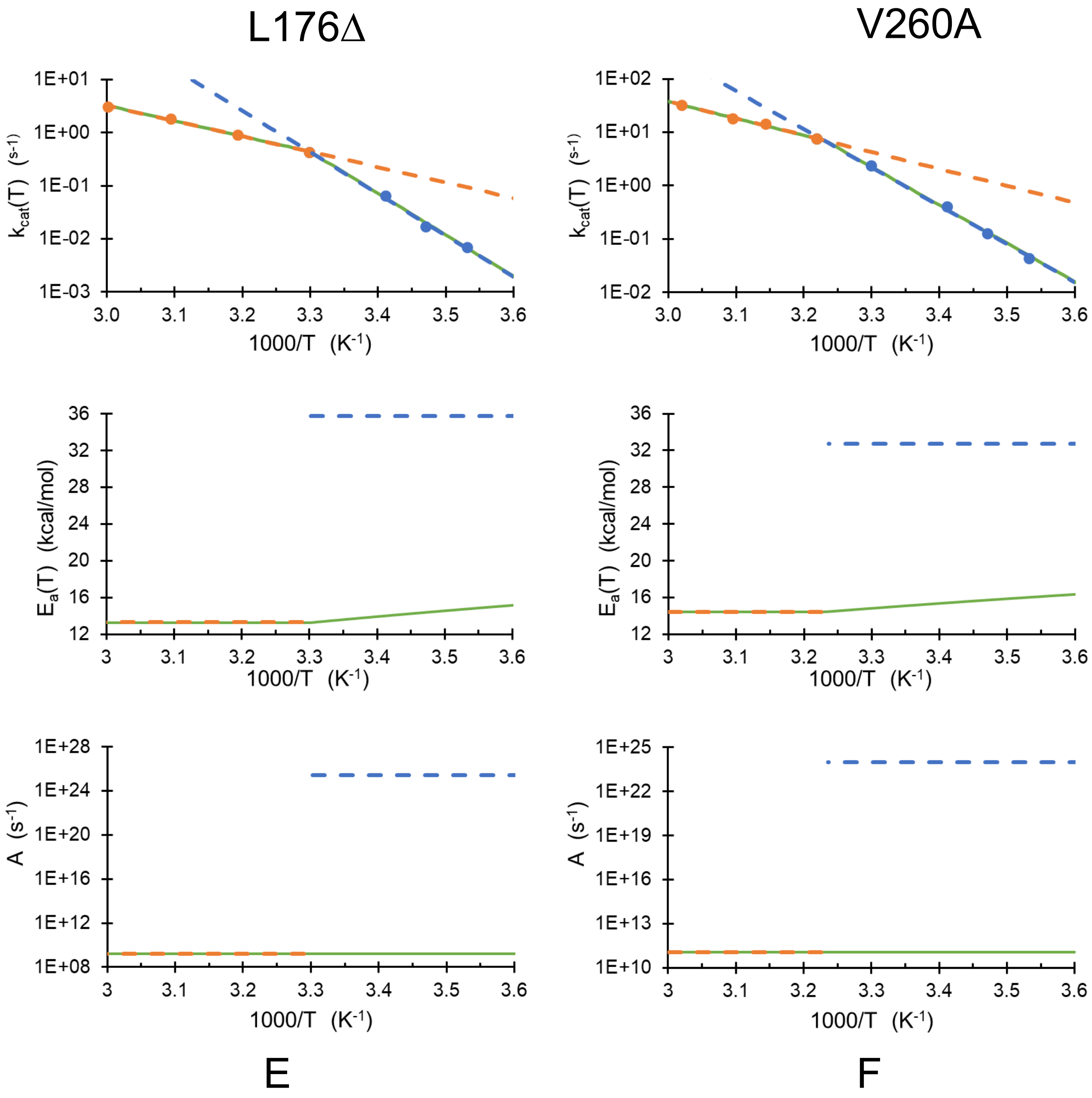


**Figure S4. (Top)** Arrhenius plot of $k_{cat}(T)$ data from (Nagel et al., 2011) for WT and 5 mutant forms of a thermophilic alcohol dehdrogenase (ht-ADH). The orange and blue dashed lines are standard Arrhenius fits to the data above and below the break temperature, respectively**. (Middle and Bottom)** The corresponding $E_a$ and $A$ values above and below the break, determined by the slope and intercept of each fit, are shown by orange and blue dashed lines. The solid green lines are fits to the data (top) and the corresponding $E_a(T)$ and $A(T)$ (middle and bottom) for each enzyme, assuming that $E_a$ is constant above the break and varies linearly with temperature below the break and that $A$ has the same value above and below the break. In all cases, the linear temperature variation of $E_a(T)$ required to replicate the observed data is much smaller than jump in $E_a$ required at the break if $E_a$ is assumed to be temperature independent away from the break.

### Tables

**Table S1.** Effect of linear variation of $E_a$ with temperature on apparent activation energy $E_a^{app}$ (given by Eq. 7a) and apparent prefactor $A^{app}$ (given by Eq. 7b) deduced from the slope and intercept of an Arrhenius plot. $E_a^{app}$ is calculated using Eq. 7a assuming a linear variation of $E_a$ about $T_0$=303 K, for several values of the temperature coefficient *a*.

| *a* (% / °C) | $\Delta E_a / E_a(T_0)$ between 273 and 333 K | $E_a^{app} / E_a(T_0)$ | $A^{app} / A$ for $E_a(T_0) = 10$ kcal/mol |
|---|---|---|---|
| -0.1 | 0.06 | 1.3 | $1.4 \times 10^2$ |
| -0.2 | 0.12 | 1.6 | $2.4 \times 10^4$ |
| - 0.5 | 0.30 | 2.5 | $8.5 \times 10^{10}$ |
| - 1 | 0.60 | 4.0 | $7.2 \times 10^{21}$ |

**Table S2.** Parameters of Arrhenius fits to $k_{cat}$ vs $1/T$ data from WT and mutant thermophilic alcohol dehydrogenases, from Tables 2 and 3 of (Nagel et al., 2011).

| Enzyme | *T* above Arrhenius break | | | *T* below Arrhenius break | | |
|---|---|---|---|---|---|---|
| | $\Delta H^{rxn,app}$ (kcal/mol) | $T\Delta S^{rxn,app}$ (kcal/mol) | $\log\left(A^{app}\right)$ | $\Delta H^{rxn,app}$ (kcal/mol) | $T\Delta S^{rxn,app}$ (kcal/mol) | $\log\left(A^{app}\right)$ |
| WT | 14.7 | -1.2 | 12.4 | 21.4 | 5.6 | 17.2 |
| L176V | 17.9 | 0.6 | 13.7 | 26.9 | 9.6 | 20.2 |
| L176A | 15.3 | 1.8 | 12.0 | 25.8 | 8.8 | 19.6 |
| L176G | 15.0 | -1.5 | 12.1 | 26.7 | 10.2 | 20.6 |
| L176D | 12.8 | -5.3 | 9.4 | 34.8 | 16.5 | 25.2 |
| V260A | 13.6 | -3.3 | 10.9 | 32.3 | 15.4 | 24.1 |

**Table S3.** Parameters of conformational sampling fit Eq. S2 to $k_{cat}/T$ vs $1/T$ data from WT and mutant thermophilic alcohol dehydrogenases, from Table 4 of (Nagel et al., 2011).

| Enzyme | *T* below Arrhenius break | | |
|---|---|---|---|
| | $\Delta H_c$ (kcal/mol) | $T\Delta S_c$ (kcal/mol) | $\Delta G_c$ (kcal/mol) |
| WT | 6.7 | 6.8 | -0.1 |
| L176V | 8.9 | 9.0 | -0.1 |
| L176A | 10.5 | 10.6 | -0.1 |
| L176G | 11.7 | 11.7 | 0 |
| L176D | 22.0 | 21.9 | 0.1 |
| V260A | 18.7 | 18.8 | -0.1 |

**Table S4.** Temperature coefficient *a* of the activation energy $E_a\left(T\right)=E_{a,0}\left(1+\alpha\left(T-T_0\right)\right)$ required to account for the large increase in apparent activation energy $E_a^{app}=E_{a,0}\left[1-\alpha T_0\right]$ and prefactor $A^{app}=A\exp\left(-\frac{\alpha E_{a,0}}{k_B}\right)$ observed below the breakpoint temperature (40 °C for V260A, 30 °C for all other variants) in **Table S2**, assuming that the underlying activation energy is constant above the breakpoint and varies linearly with temperature below the breakpoint. All values given here were obtained by refitting the raw $k_{cat}(T)$ data in Refs. 15 and (Kohen, Amnon, Cannio, Raffaele, & Bartolucci, Simonetta, 1999).

| Enzyme | T above | *T* below Arrhenius break | | | |
|---|---|---|---|---|---|
| | $E_a$ (kcal/mol) | $\alpha$ (%/°C) | Change in $E_a(T)$ between 0 and 30°C (kcal/mol) | $E_a^{app}$ (kcal/mol) | $\log\left(A^{app}\right)$ |
| WT | 15.5 | -0.14 | -0.65 | 22.1 | 17.7 |
| L176V | 18.5 | -0.17 | -0.96 | 28.2 | 20.6 |
| L176A | 15.8 | -0.22 | -1.04 | 26.3 | 19.5 |
| L176G | 16.6 | -0.21 | -1.02 | 26.9 | 20.3 |
| L176D | 13.3 | -0.56 | -2.22 | 35.7 | 25.5 |
| V260A | 14.4 | -0.41 | -1.78 | 33.0 | 24.0 |

**Table S5.** Temperature coefficient a of $\Delta G^{rxn}\left(T\right)=\Delta G^{rxn}\left(T_{break}\right)\left[1+\alpha'\left(T-T_{break}\right)\right]$ required to fit the observed E-P plot slope and intercept and corresponding values of $\Delta H^{rxn,app}$ and $\Delta S^{rxn,app}$ observed at temperatures below the breakpoint $T_{break}$, assuming that $\Delta H^{rxn}$ and $\Delta S^{rxn}$ above the break are constant and so can be obtained directly using the standard E-P fit. Rather than using values for $\Delta H^{rxn,app}$ and $T\Delta S^{rxn,app}$ in Table S2 (where the $T$ value was not specified in Ref. (Nagel et al., 2011)), the values were obtained by fitting the raw $k_{cat}(T)/T$ data. $\Delta G_0^{rxn}$ is evaluated at the breakpoint temperature.

| Enzyme | $T$ above Arrhenius break | | | | $T$ below Arrhenius break | | |
|---|---|---|---|---|---|---|---|
| | $\Delta H^{rxn}$ (kcal/mol) | $\Delta S^{rxn}$ (kcal/mol) | $\Delta G_0^{rxn}$ (kcal/mol) | $\alpha_0'$ (%/°C) | $\Delta H^{rxn,app}$ (kcal/mol) | $\Delta S^{rxn,app}$ (kcal/mol) | $\alpha'$ (%/°C) |
| WT | 14.8 | $-2.2\times10^{-3}$ | 15.5 | 0.014 | 21.5 | $20.1\times10^{-3}$ | -0.13 |
| L176V | 17.8 | $0.9\times10^{-3}$ | 17.6 | -0.005 | 27.5 | $33.4\times10^{-3}$ | -0.19 |
| L176A | 15.2 | $-6.7\times10^{-3}$ | 17.2 | 0.039 | 25.7 | $27.5\times10^{-3}$ | -0.16 |
| L176G | 16.0 | $-2.4\times10^{-3}$ | 16.7 | 0.014 | 28.1 | $38.0\times10^{-3}$ | -0.22 |
| L176D | 12.6 | $-19.0\times10^{-3}$ | 18.4 | 0.010 | 35.2 | $54.9\times10^{-3}$ | -0.30 |
| V260A | 13.8 | $-10.7\times10^{-3}$ | 17.1 | 0.062 | 32.2 | $48.5\times10^{-3}$ | -0.29 |
